# Heterogeneous *pdgfrβ+* cells regulate coronary vessel development and revascularization during heart regeneration

**DOI:** 10.1101/2021.04.27.441161

**Authors:** Subir Kapuria, Haipeng Bai, Juancarlos Fierros, Ying Huang, Feiyang Ma, Tyler Yoshida, Antonio Aguayo, Fatma Kok, Katie M. Wiens, Joycelyn K. Yip, Megan L. McCain, Matteo Pellegrini, Mikiko Nagashima, Peter F. Hitchcock, Nathan D. Lawson, Michael MR Harrison, Ching-Ling Lien

## Abstract

Endothelial cells emerge from the atrioventricular canal (AVC) to form nascent coronary blood vessels in the juvenile zebrafish heart. We found that *pdgfrβ* is first expressed in the epicardium around the AVC and later becomes localized mainly in the mural cells. *pdgfrβ* mutant fish display severe defects in mural cell recruitment and coronary vessel development. *pdgfrβ+* mural cells are heterogeneous and those associated with coronary arteries also express *cxcl12b*. Mural cells positive for both *pdgfrβ* and *cxcl12b* transgenic reporters had elevated expression of smooth muscle cell genes. Interestingly, these mural cells were associated with coronary arteries even in the absence of Pdgfrβ, although smooth muscle gene expression was downregulated in these cells. We found that *pdgfrβ* expression dynamically changes in the epicardium derived cells, which we found to be a heterogeneous population. *mdka* was identified as a gene upregulated in subpopulations of *pdgfrβ*+ cells during heart regeneration. However, *pdgfrβ* but not *mdka* mutants showed defects in heart regeneration. Our results demonstrated that *pdgfrβ+* cells and Pdgfrβ signaling are essential for coronary development and heart regeneration.

**SUMMARY STATEMENT:** Heterogeneous *pdgfrβ* positive cells are present in developing and regenerating zebrafish hearts and are required for development of mural cells and their association with the nascent coronary vessels during zebrafish heart development and regeneration.

## INTRODUCTION

Coronary heart disease is the leading cause of human mortality worldwide. Understanding the mechanisms of coronary vessel development and revascularization has thus drawn much attention in hope to advance the treatment of heart disease. In contrast to mammals, zebrafish is a well-established, genetically tractable model organism that shows remarkable regenerative capacity in the adult heart. Fast vascularization is essential to regenerate damaged zebrafish hearts (Marin-Juez et al., 2016). The presence of well-structured coronary vasculature and ease of imaging the developing and adult hearts make zebrafish an ideal system for exploring cellular and molecular mechanisms of coronary vessel formation. We previously reported that the Cxcr4a-Cxcl12b chemokine axis guides newly emerged endothelial sprouts from the atrioventricular canal (AVC) to undergo angiogenesis and gradually cover the juvenile heart ventricle during development (Harrison et al., 2015). The cellular and molecular mechanisms that govern the maturation and maintenance of the developed coronary network remain unclear.

Mural cells (including both pericytes and smooth muscle cells) are a collection of diverse supporting cells that are recruited onto endothelial cells and cover the circulatory vessels as single or multiple cell layers. They regulate vessel development, stability, and physical functions such as vessel contractility (Armulik et al., 2011). Platelet-derived Growth Factor b and its receptor *β* (Pdgf-b/Pdgfrβ) regulate mural cell recruitment onto endothelial cells (Armulik et al., 2005, Armulik et al., 2011, Hellstrom et al., 1999, Lindahl et al., 1997, Lindblom et al., 2003, Winkler et al., 2010). Endothelial cells express Pdgf-b and mural cells express the receptor, Pdgfrβ (Winkler et al., 2010). Genetic disruption of *Pdgf-b* or *Pdgfrβ* significantly decreases mural cell coverage in vessels throughout the mouse embryo (Hellstrom et al., 1999). In the central nervous system (CNS), loss of mural cell coverage in the vasculature makes the blood vessels hyperplastic (significantly more endothelial cells per vessel), dilated, and susceptible to hemorrhage (Lindahl et al., 1997).

In mouse hearts, mural cells originate from epicardial (Cai et al., 2008, Mellgren et al., 2008) and endocardial cells (Chen et al., 2016). During development, pericytes are recruited to micro-vessels and mature further to become smooth muscle cells on coronary arteries. Smooth muscle cell differentiation is promoted with the commencement of blood flow through the arterial vessel that in turn stimulates Notch activation in the pericytes (Volz et al., 2015). PDGFRβ signaling is essential for coronary smooth muscle development in both mouse and avian hearts (Mellgren et al., 2008, Smith et al., 2011, Van Den Akker et al., 2005, Van Den Akker et al., 2008) and pericytes are decreased in *Pdgfrβ* null mice (Volz et al., 2015). It is not yet clear how diverse the cardiac mural cells populations are and how PDGFRβ signaling regulates different mural cell populations.

Herein we describe the *pdgfrβ* expression patterns, the epicardial origin of mural cells, and heterogeneity of *pdgfrβ+* mural cells during zebrafish heart coronary vessel development. We found a subpopulation of *pdgfrβ+* mural cells associated with coronary arteries also expresses *cxcl12b*. These *pdgfrβ*; *cxcl12b* double positive mural cells express makers of smooth muscle cells. Interestingly, these mural cells positive for both *pdgfrβ* and *cxcl12b* transgenic reporters still associated with the coronary arteries even though other *pdgfrβ*+ mural cells detached from non-arterial coronary vessels in the absence of Pdgfrβ. However, single cell RNA sequencing (scRNAseq) analyses revealed downregulation of smooth muscle gene expression in these mural cells associated with coronary arteries in *pdgfrβ* mutants. Using a novel fluidic device-based culture and live imaging system, we further demonstrated that *pdgfrβ* expression dynamically changes in the epicardium derived cells (EDPCs) and that pre-existing mural cells migrate with angiogenic endothelial cells during heart regeneration. Using scRNAseq, we found distinct populations of *pdgfrβ+* EDPCs and pre-existing mural cells in regenerating zebrafish hearts. Furthermore, *pdgfrβ* mutants showed defects in mural cell association with coronary endothelial cells during heart regeneration; this severely compromised the regenerative response. We identified *mdka* as a candidate gene that showed similar trend in gene expression patterns in EDPCs as *pdgfrβ*. However, we did not observe defects in heart regeneration in *mdka* mutants. Our results suggest that heterogeneous *pdgfrβ+* cells and Pdgfrβ signaling are essential for coronary development and heart regeneration.

## RESULTS

### *pdgfrβ* expression and the origins of *pdgfrβ* positive mural cells in developing zebrafish heart

To characterize *pdgfrβ* spatiotemporal expression patterns in the developing zebrafish heart, we utilized a *pdgfrβ*:*mCitrine* transgenic reporter (Vanhollebeke et al., 2015). In juvenile fish, the formation of coronary endothelial cells initiated on the ventricle around 36-49 days post fertilization (dpf) (∼14-18 mm in body length) [marked by *Tg(fli1a: DsRed)* in Fig. 1A’’]. Interestingly, *pdgfrβ* expression was consistently observed in the bulbus arteriosus (BA) and around the atrioventricular canal (AVC) at the late larval stage (24 dpf), before any coronary endothelial cells emerged (Fig. 1A). In early juvenile fish, *pdgfrβ* expression spread along the AVC at 31 and 43 dpf. At the late juvenile stage by 55-74 dpf, *pdgfrβ* expression became localized in the mural cells accompanying the nascent coronary endothelial cells sprouting out from the AVC (Fig. 1A’”, Fig. S1A).

**Figure 1.**
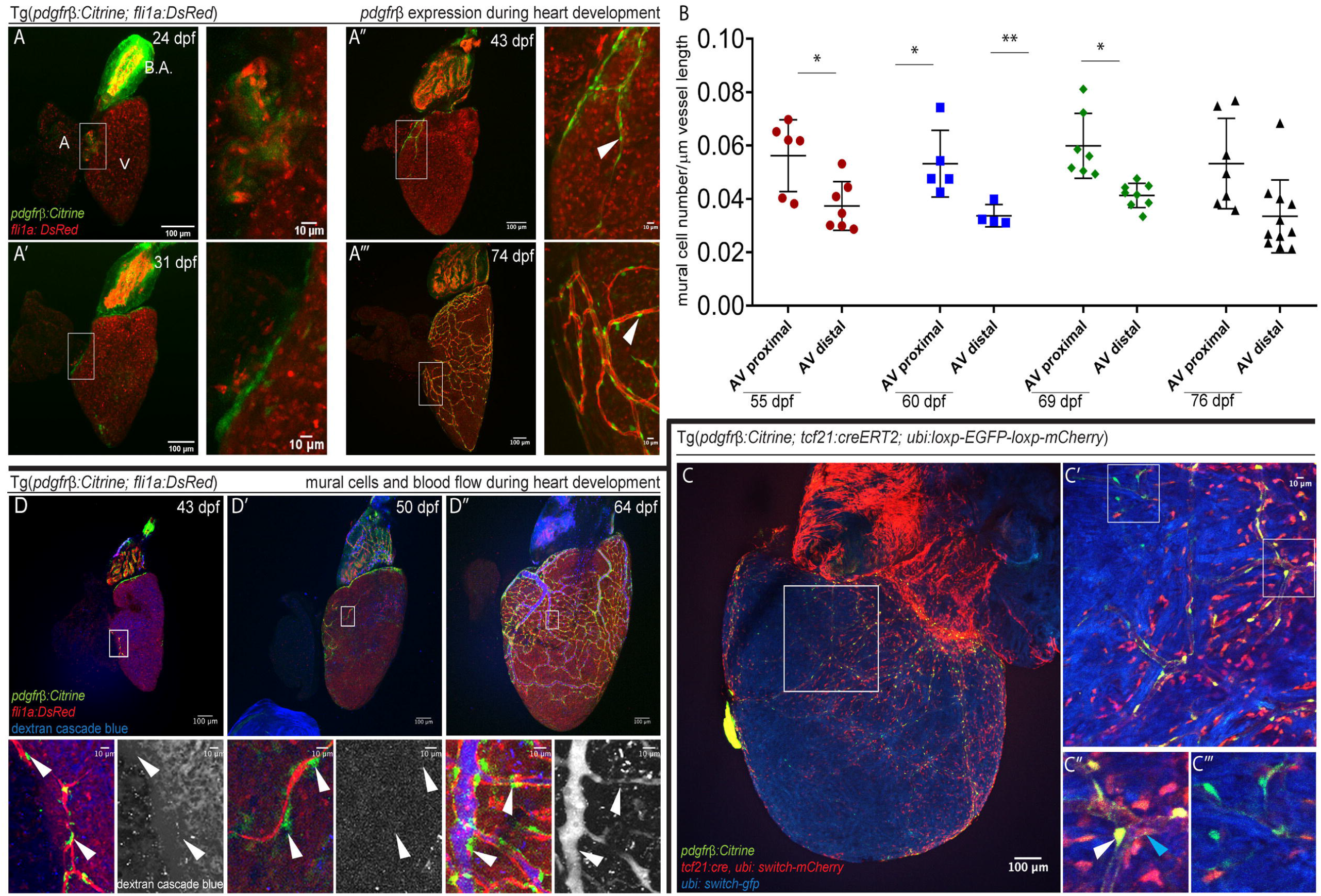
*pdgfrβ* expression and mural cell origin in the developing zebrafish heart. (A-A’”) *pdgfrβ* expression during heart development at 24 dpf (A), 31 dpf (A’), 43 dpf (A’’), and 74 dpf (A’’’). Zoomed-in images on the right. V, ventricle; A, atrium; B.A. bulbus arteriosus. n = 5 hearts for each time point. (B) Quantification of *pdgfrβ+* mural cell association. Mural cell number per μm of the coronary vessels proximal or distal to AVC at different timepoints (55, 60, 69, and 76) dpf. Boxes in (A) indicate the vessels quantified as proximal to (near) AVC. n = 5-7 hearts for each time point. Error bars, standard deviation of mean. T-test (*p<0.05,**p < 0.01). (C-C”‘) *tcf21+* lineage traced cells contribute to *pdgfrβ+* mural cells. (C) Representative image of *tcf21* lineage traced fish at 87 dpf. *ubi:switch-GFP* (artificially colored blue) switches to *ubi:switch-mCherry* after recombination. *pdgfrβ*:*Citrine*, green. (C’-C”‘) Zoomed-in images. (C’’) *tcf21* lineage-traced *(mCherry)* cells co-labelled with *pdgfrβ+* cells (white arrow) and *pdgfrβ-*, non-mural cell (blue arrow). C”‘, *pdgfrβ+* cells (green) that are negative for *tcf21* lineage. n=3. (D-D”) Mural cell development around the coronary vessels does not depend on blood flow. Coronary angiography (Dextran, blue) performed to monitor blood flow at 43 (D), 50 (D’) and 64 (D’’) dpf. White arrowhead indicates *pdgfrβ+* mural cells in the zoomed-in images. n = 5-6 hearts for each time point.

As growing coronary vessels covered the ventricle, mural cells remained associated with large and small coronary vessels (Fig. 1A’’’). Throughout development, mural cell density remained greater on the coronary vessels closer to the AVC (defined as the proximal area to AVC) than the growing end of the vessels (regions distal to the AVC) (Fig.1B, S1A). The co-localization of *pdgfrβ:mCitrine* and *tcf21:DsRed* (which marks epicardium) around the AVC at 33 dpf (Fig. S1B) suggests that the origins of *pdgfrβ+* cells might be the epicardium. To confirm this, we performed lineage tracing experiments using the *tcf21:CreERT2* line combined with *pdgfrβ:Citrine* and the *ubi:Switch* reporter. We observed that 76.15% of *pdgfrβ+* cells were co-labeled with mCherry, indicating most *pdgfrβ+* cells were derived from *tcf21+* epicardial lineage (Fig. 1C-C’’, white arrow in C’’, Fig. S1C). 23.85% of mural cells (Fig. S1C) were not co-labeled with mCherry, suggesting that either *CreER-loxP* recombination was not efficient in these cells. Alternatively, these *pdgfrβ+* cells were derived from other sources (Fig. 1C’’’). Furthermore, we observed *tcf21:CreERT2* labeled cells that did not express *pdgfrβ* (Fig. 1C’’, blue arrow), consistent with the notion that epicardium can also contribute to other cell types such as fibroblasts in zebrafish (Sanchez-Iranzo et al., 2018) and Pdgfrβ+ pericyte progenitors in addition to fibroblasts in mice (Ivey et al., 2018, Volz et al., 2015).

Next, we investigated if the appearance of *pdgfrβ*-expressing mural cells depended on blood circulation through the developing coronary vessels. We performed coronary angiography using *Tg(pdgfβ:Citrine; fli1a:DsRed)* juvenile zebrafish. *pdgfβ:Citrine+* mural cells were observed at 43 dpf on nascent coronary vessel sprouts in the absence of blood flow (Fig. 1D). By 50 dpf, initiation of blood flow through major vessels was observed in some hearts. However, in these juvenile fish *pdgfβ:Citrine+* mural cells were already attached to the immature vessel plexus without any detectable blood circulation (Fig. 1D’). All major vessels had blood flow by 64 dpf (Fig. 1D’’). These results indicated that blood flow is not necessary for *pdgfβ:Citrine+* mural cell association with coronary vessels.

### Pdgfrβ regulates mural cell recruitment to the coronary vessels and is required for coronary vessel development

To determine the role of Pdgfrβ during coronary vessel development, we examined zebrafish carrying the *pdgfrβ*^*um148*^loss-of-function allele [*pdgfrβ*^*-/-*^, (Kok et al., 2015)]. Consistent with the findings in Pdgfrβ knockout mice (Lindahl et al., 1997), *pdgfrβ*^*-/-*^ fish displayed hemorrhage in the brain (data not shown), and the adult brain vasculature became significantly dilated with reduced branching density and decreased mural cell association (Fig. S2A). We did not observe hemorrhage in the heart ventricle (data not shown). In *pdgfrβ*^*-/-*^and *pdgfrβ* heterozygous (*pdgfrβ*^*+/-*^) mutants, coronary vessels developed although not as efficiently as *pdgfrβ*^*+/+*^fish. Both *pdgfrβ*^*-/-*^ and *pdgfrβ*^*+/-*^ ventricles had reduced coverage of coronary vessels at 98 dpf (Fig. 2A, S2B’). Unlike in *pdgfrβ*^*+/-*^ fish, where coronary vessel development was delayed, these defects in coronary vessel coverage permanently reduced by 167 dpf in *pdgfrβ*^*-/-*^ homozygous mutants (herein we use *pdgfrβ* mutants) (Fig. 2A, A’). Isolated endothelial cells, which fail to form continuous vessel networks, were observed in *pdgfrβ* mutants (Fig. 2A, S2B, marked by asterisks). Mural cell association with small vessels decreased significantly, but association with some of the large vessels remained unaffected in the *pdgfrβ* mutants (Fig. 2B, B’, S2C). Further analysis revealed that among all large vessels, narrow large vessels [coronary artery (Harrison et al., 2015)] maintained *pdgfrβ+* mural cell association (Fig. 2B, S2C white arrowhead), while the wide large vessels (vein-like) significantly lost *pdgfrβ+* mural cells (Fig. 2B, S2C, yellow arrowhead). Interestingly, in contrast to brain vessels that became dilated (Fig. S2A), the diameter of both wide large vessels and coronary arteries significantly decreased in *pdgfrβ* mutants (Fig. S2D), suggesting a potential organ-specific mechanism of *pdgfrβ*+ mural cell regulating blood vessel formation/maturation. Taken together, these data suggest that different mechanisms might be utilized to regulate mural cell recruitment or maintenance along different subtypes of coronary vessels.

**Figure 2.**
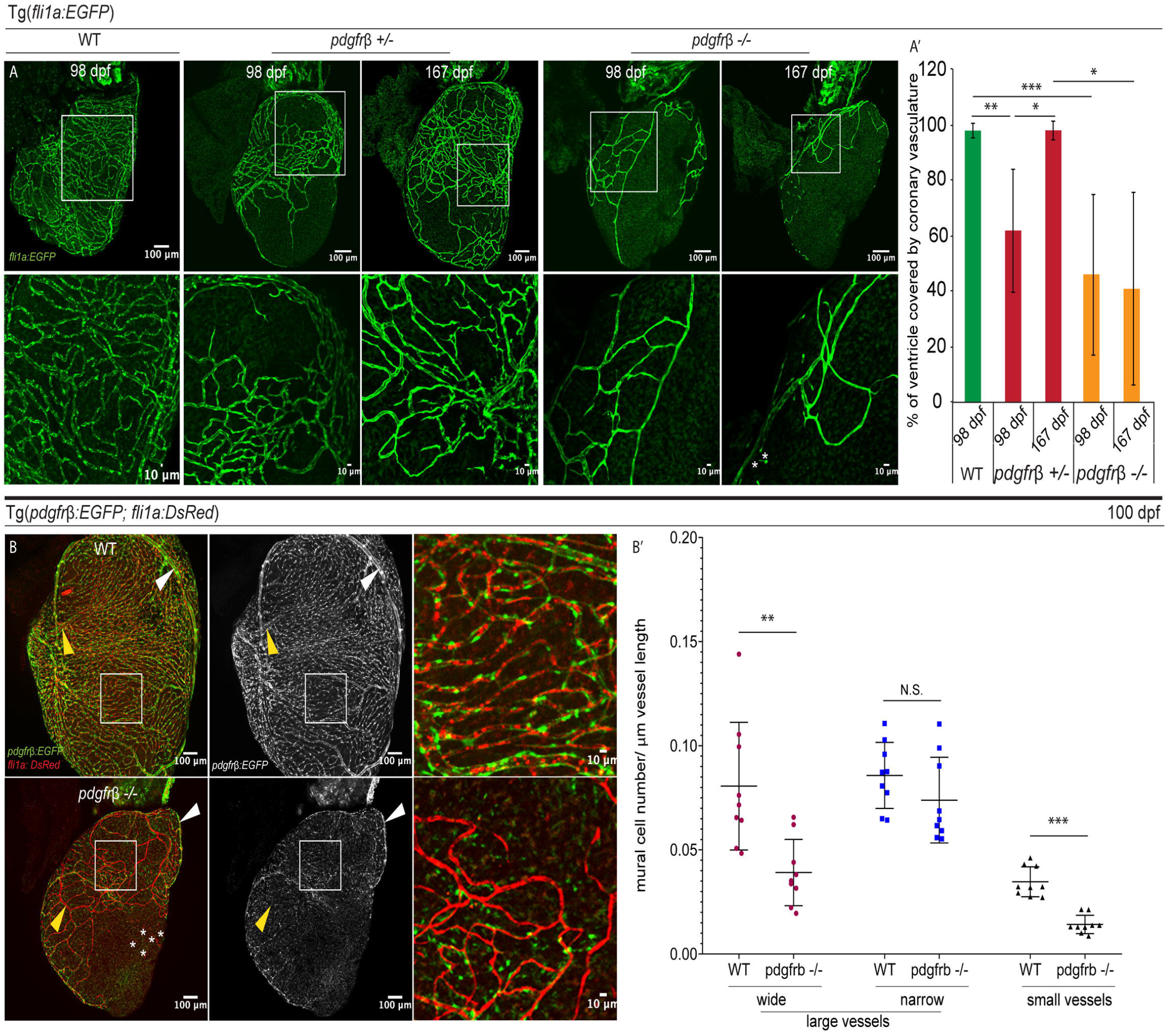
Pdgfrβ regulates mural cell number, association, and development of the coronary vessels. (A-A’) Coronary vessel coverage of the ventricle. (A) Imaging of *fli1a:*EGFP in WT (*pdgfrβ*^+/+^), heterozygous (*pdgfrβ*^+/-^), and homozygous mutant (*pdgfrβ*^-/-^) fish at 98 and 167 dpf. * indicates isolated endothelial cells. (A’). Quantification of coronary vessel coverage as the percentage of the ventricle area. *Pdgfrβ*^+/-^ fish have decreased vessel coverage at 98 dpf (∼61.8%, n = 6 fish) but recovered by 167 dpf (∼97.85%, n = 4 fish). *Pdgfrβ*^-/-^ mutant have decreased vessel coverage at both 98 dpf (∼41.45%, n = 8 fish) and 167 dpf (∼40.85%, n = 4 fish), WT (97.76%, n=7). Error bars, standard deviation of mean. One-way ANOVA (*p < 0.05, **p < 0.01, ***p < 0.001). (B-B’) Mural cell association with different types of coronary vessels affected differently in *pdgfrβ*^-/-^ mutant. (B) Mural cell [*Tg*(*pdgfrβ:*EGFP)] association with the small coronary vessels/capillaries [*Tg*(*fli1a:DsRed*)] decreased at 100 dpf in *pdgfrβ*^-/-^ mutant heart. Mural cells are maintained around large narrow (artery like) vessels (white arrow), but the wider large (vein like) vessels (yellow arrow) lack mural cells. * indicates isolated endothelial cells. (B’) Quantification of the mural cell recruitment on large and small coronary vessels. Mural cell number/μm vessel length on the narrow large vessels is not significantly different between *pdgfrβ*^*+/+*^ and *pdgfrβ*^*-/-*^. The mural cell recruitment on the wide large vessels and small coronary vessels is significantly decreased in *pdgfrβ*^*-/-*^ heart ventricles. n = 3 vessels x 3 - 5 heart ventricle of *pdgfrβ*^*+/+*^ and *pdgfrβ*^*-/-*^fish. Error bars, standard deviation of mean. T-test (***p < 0.001, **p < 0.01, N.S. = not significant > 0.05).

### *pdgfrβ; cxcl12b* expressing cells covering the coronary arteries are smooth muscle like mural cells that maintain endothelial cell association in *pdgfrβ*^-/-^ fish

We reported that epicardium-derived *cxcl12b:Citrine*-expressing mural cells surrounded *cxcr4a+* arterial endothelial cells (Harrison et al., 2015). We also observed *pdgfrβ+* cells covering coronary arteries (Fig. 2B, S2C). Therefore, we examined whether *cxcl12b:Citrine+* mural cells also expressed *pdgfrβ:EGFP*. We observed that 82.39% of the mural cells on the coronary arteries expressed both *cxcl12b* and *pdgfrβ* reporters (Fig. 3A’-A’’’ blue arrow, 3A””). *pdgfrβ:EGFP+* only (Fig. 3A’-A’’’, green arrow) and *cxcl12b:mCitrine+* only (Fig. 3A’-3A’’’, red arrow) mural cells were less abundant on coronary arteries (12.88% and 4.73% respectively) (Fig. 3A””).

**Figure 3.**
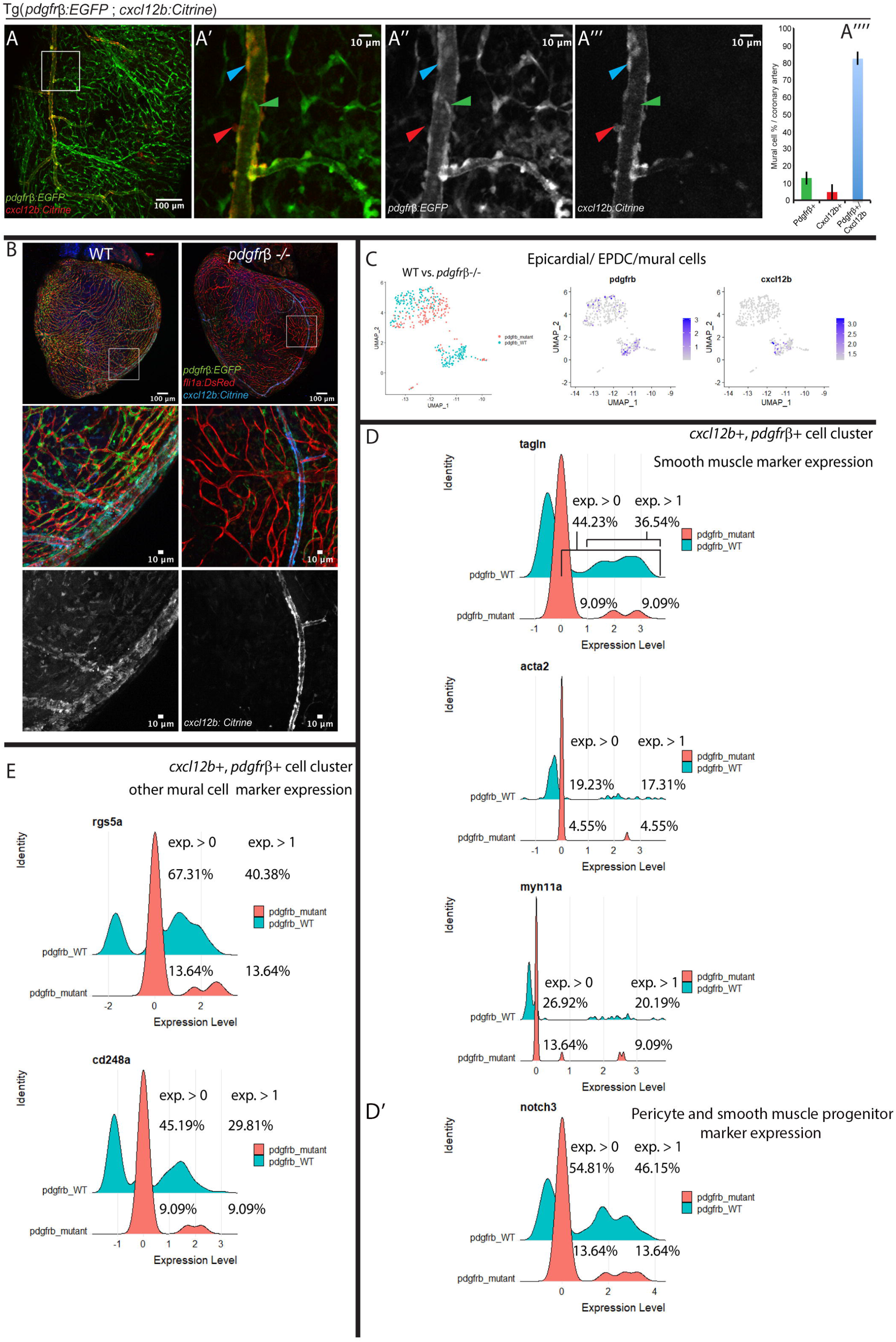
*pdgfrβ ;cxcl12b*+ arterial mural cells are smooth muscle like cells and their association with endothelial cells is *pdgfrβ* independent. (A-A’’’’) Majority of the *pdgfrβ*+ mural cells express *cxcl12b* on the large narrow vessels (coronary artery). (A-A’’’). On the main trunk of the coronary artery, most *pdgfrβ*:*EGFP*+ mural cells (green) express *cxcl12b:Citrine* (blue arrow). A few mural cells only express *pdgfrβ* (green arrow) or *cxcl12b* (red arrow). *cxcl12b* expression in the *pdgfrβ*+ mural cells gradually decrease on the branches away from the main coronary artery (A’’’). (A’’’’) Quantification of *pdgfrβ*+;*cxcl12b*+ (∼82.4% of the mural cells on the coronary artery), *pdgfrβ*+ only (∼12.9%) and *cxcl12b*+ only (∼4.7%) mural cells (n = 5 hearts, 1-2 coronary arteries in each heart, 20X confocal images used for quantification). (B) In *pdgfrβ*^-/-^, *pdgfrβ:EGFP* and *cxcl12b:Citrine* expressing mural cells remain associated with coronary artery while other vessels lose mural cell association. (C) FACS isolated *pdgfrβ:EGFP*+ EPDCs and mural cells form two distinct clusters in the UMAP plot where cells in the both clusters express *pdgfrβ*, only one cluster cells express *cxcl12b*. Cells from the wildtype fish and *pdgfrβ*^-/-^ fish are distinct from each other in the same clusters in the UMAP. (D-E) The ridge-plot of different marker gene expression in the *cxcl12b*+;*pdgfrβ*+ cell cluster of wild type vs. *pdgfrβ*^-/-^ mutant fish. The percentage of cells with expression level > 0 (exp. > 0) and >1 (exp. > 1) is indicated for each ridge-plot. (D-D’) The ridge plots for the smooth muscle marker genes (*acta2, myh11a, tagln*) and pericyte and smooth muscle progenitor marker, *notch3*. All the marker gene expressions decrease in *pdgfrβ*^-/-^ cells compared to wild type cells. (E) Other mural cell markers (rgs5a, cd248a) decrease in the *pdgfrβ*^-/-^ cells.

We examined whether these *cxcl12b+* cells along the coronary artery are affected in *pdgfrβ* mutants. Consistent with the finding using the *pdgfβ:EGFP* reporter (Fig. 2B, Fig. S2C), we found that *cxcl12b:Citrine+* (also *pdgfrβ*:*EGFP*+) mural cells remained associated with coronary arteries whereas other coronary vessels lose mural cells and the diameter of the coronary artery is significantly decreased in *pdgfrβ* mutants (Fig. 3B). These results suggest that *pdgfrβ*+ mural cells are heterogeneous on the heart ventricle; some of them are *pdgfrβ* only, and some of them are double positive with *cxcl12b*. To further examine these *pdgfrβ*:*EGFP*+ and *cxcl12b:Citrine*+ double positive (*pdgfrβ*+;*cxcl12b*+) mural cells and other *pdgfrβ*:*EGFP*+ only cells in controls vs. *pdgfrβ* mutants, we performed single cell RNA sequencing (scRNAseq). We FACS sorted *pdgfrβ:EGFP+* cells from *pdgfrβ* mutants and control fish to enrich the mural cells although different cell types (especially cardiomyocytes) were also collected, likely due to the autofluorescence (Fig. S3.1A, B’). In Uniform Manifold Approximation and Projection (UMAP) plot, the *pdgfrβ*+ cells form two distinct clusters and the wild type cells and *pdgfrβ* mutant cells took distinct position in each cluster, reflecting their overall gene-expression differences (Fig. 3C). *pdgfrβ*+ cells are mainly in clusters 3 and 6; so are EGFP transcripts+ cells while *cxcl12b*+ cells are specifically in cluster 6. (Fig. S3.1A’, S3.1B). The epicardial markers, *tcf21, tbx18* are specifically found in cluster 3, suggesting this cluster consists of epicardium and epicardial derived cells (EPDCs) (Fig. S3.1B). Cluster 6 is likely representing the *cxcl12b*+, *pdgfrβ*+ arterial mural cells (Fig. 3C).

Interestingly, compared to *pdgfrβ*+ only cell-cluster, more *pdgfrβ*+;*cxcl12b*+ cluster cells showed higher expression of different vascular smooth muscle cell markers (*acta2, myh11a, tagln*) (Fig. 3D, S3.2A). Furthermore, more cells in *pdgfrβ*+;*cxcl12b*+ cluster express higher *notch3* (expression level > 1 in the ridge plots) which regulates vascular pericytes differentiation into smooth muscle cells (Volz et al., 2015) (Fig. 3D’, S3.2A’). Compared to wildtype controls, *pdgfrβ+;cxcl12b*+ mural cells show reduced *notch3* and other smooth muscle cell genes expressions in *pdgfrβ* mutants, indicating impaired smooth muscle cell differentiation even though they are associated with coronary artetries (Fig. 3D-D’). Among other established mural cell markers, *rgs5a, cd248a*, and *kcne4* (He et al., 2016, Whitesell et al., 2019) that showed prominent expression among these clusters, *rgs5a, cd248a* express more in the *pdgfrβ*+; *cxcl12b*+ cluster (Fig. 3E) whereas *kcne4* is expressed in both clusters (Fig. S3.2C). All these mural cell markers showed reduced expression in *pdgfrβ* mutant cells in both clusters, indicating overall reduced mural cell identity or differentiation (Fig. 3E, S3.2C). Few other mural cell markers, *abcc9, desma* (He et al., 2016) showed expression in the *pdgfrβ*+;*cxcl12b*+ cluster and *cspg4, anpepa* (He et al., 2016) expressed in a few cells of the *pdgfrβ*+ only cell-cluster (data not shown) indicating overall heterogeneity of the coronary mural cell population. Among epicardial markers, *tbx18* and *tcf21* showed prominent expression in the *pdgfrβ*+ only cell-cluster and much less expression in the *pdgfrβ*+;*cxcl12b*+ cluster (Fig.S3.2B). *pdgfrβ*^*-/-*^ cells showed reduced expression of *tcf21* and *tbx18* in the *pdgfrβ*+;*cxcl12b*+ cluster. However, in the *pdgfrβ*+ only cell-cluster the expression is less but comparable to wild type (Fig. S3.2B). These analysis results suggested that *pdgfrβ*+ cells are derived from epicardium and cells in *pdgfrβ*+;*cxcl12b*+ cluster are smooth muscle cell-like. Furthermore, cells in these 2 clusters are affected differently in *pdgfrβ* mutants.

The *pdgfrβ*+ only cell-cluster is further analyzed to determine the effect of *pdgfrβ*^*-/-*^ mutation on these cells. This cluster was isolated and re-clustered into 5 sub-clusters to explore the internal gene expression differences among the cells in the cluster. Among these sub-clusters, sub-cluster 0 and 2 are more represented by *pdgfrβ*^*-/-*^ cells, and subcluster 1 and 3 are more represented by wild type cells. Gene Ontology (GO) term analysis was performed to predict cellular functions. *pdgfrβ*^*-/-*^ specific sub-clusters are enriched with GO-terms for “extracellular region”, “regeneration”, “extracellular matrix”, “oxidoreductase activity”, “transmembrane transporter activity”. Wild type specific sub-clusters are enriched with GO-terms for “blood vessel development”, “Rho protein signal transduction”, “small GTPase signal transduction”, “extracellular matrix”, “cell adhesion”, “muscle organ development” (Fig. S3.1C-C’). This GO-term analysis predicts while wildtype *pdgfrβ*+ only cells regulate angiogenesis, structural integrity maintenance, neighboring cardiomyocyte development, similar cells in *pdgfrβ*^*-/-*^ condition are stressed, and show damage responsive activities even without any injury.

### *pdgfrβ* expression dynamically changes during heart regeneration

Our previous observation of *pdgfrβ* upregulation and the presence of mural cells in the regenerating area of the zebrafish heart (Kim et al., 2010) prompted us to characterize *pdgfrβ* expression patterns during regeneration. The apical regions of *Tg*(*pdgfrβ:Citrine;fli1a:DsRed*) fish were injured and imaged at different days post amputation (dpa). At 1 dpa, there were no obvious *pdgfrβ* expressing regions near the wound area except in mural cells around the pre-existing coronary vessels. Starting from 3 dpa, patchy *pdgfrβ* expression was observed around the border of the amputated area (Fig. 4A). This expression gradually expanded significantly at 7 dpa and 10 dpa, plateaued at 14 dpa, and decreased at 30 dpa (Fig. 4A, A’, S4A). The mural cells associated with pre-existing coronary endothelial cells continued to express *pdgfrβ* (Fig. S4A). This patchy *pdgfrβ* expression was likely in epicardium or EPDCs because they were also positive for the epicardial marker *tcf21* (Fig. 4B). These *pdgfrβ+* epicardial cells or EDPCs migrated and enveloped the regenerating area while the *pdgfrβ* expressing mural cells remained associated with the coronary endothelial cells migrating into the regenerating area.

**Figure 4.**
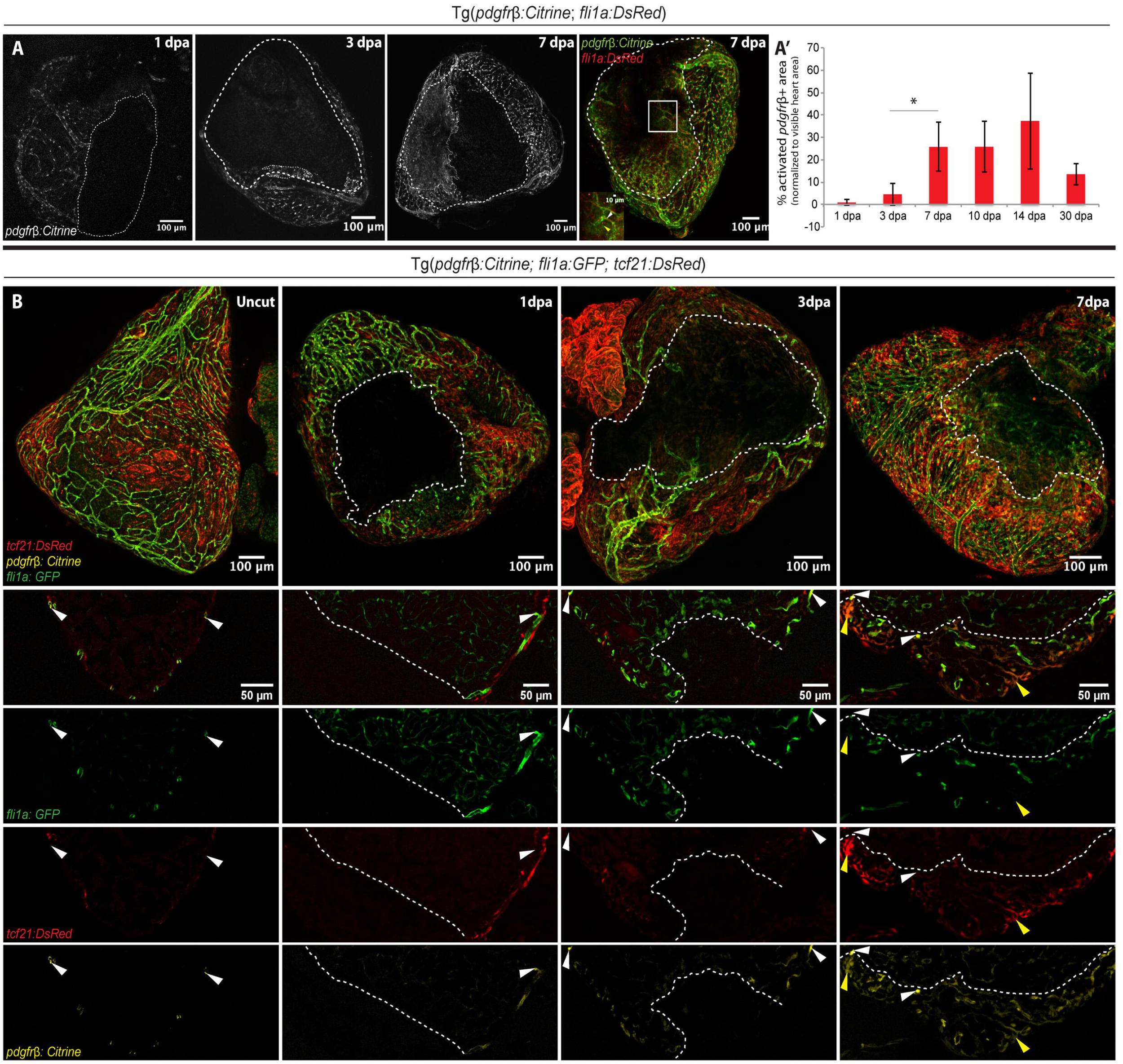
Dynamic *pdgfrβ* expression in the injured area during regeneration. (A-A’) *pdgfrβ* expression early during heart regeneration. (A) *Tg(pdgfrβ:Citrine; fli1a:DsRed)* fish hearts at 1, 3, and 7dpa. White dashed line: injured area. White dotty line: *pdgfrβ* expressing area. Inset: white arrowhead, *pdgfrβ* expression in mural cells; yellow arrowhead, non-mural cell *pdgfrb* expression. (A’) Quantification of *pdgfrβ*+ area as the percentage of the imaged heart area. Error bars, standard deviation of mean. One-way ANOVA (*p < 0.05). 1d, 3d, 30d; n = 3, 7d; n = 5, 10d, 14d; n = 4. (B) *pdgfrβ* expression in epicardium. *Tg(pdgfrβ:Citrine; fli1a:EGFP; tcf21:DsRed)* fish hearts imaged as whole-mount and sections in uncut, 1, 3, and 7 dpa. White arrowhead, *pdgfrβ* expression in mural cells; yellow arrowhead, non-mural cell *pdgfrβ* expression.

We developed a novel fluidic device-based long-term explant culture for live imaging (Yip et al., 2020) to further confirm the dynamic changes in *pdgfrβ* expression patterns and potential interactions between *pdgfrβ+* cells with other cell types,. Hearts from transgenic zebrafish *Tg*(*pdgfrβ:EGFP; fli1a:DsRed*) were injured and allowed to recover *in vivo* for 5-10 days, dissected out from the fish, then placed in the device for live imaging for 72-120 hours. *pdgfrβ:EGFP* expression at 5-6 dpa showed diffuse expression in the epicardium within most of the wound site and more localized expression in mural cells towards the edge and outside the wound site (Supplementary Movie 1). As regeneration proceeded, *pdgfrβ+* epicardial cells closed in to cover the entire wound area by 6 dpa. Following this, punctate, mural cell-like expression was observed within the regenerating region. The epicardial cells, or the diffuse expression observed within them, were highly dynamic within the wound area compared to the more stable behavior of existing mural cells (Supplementary Movie 2). These mural cells migrated into the wound site alongside the individual endothelial cell to which they were attached (Supplementary Movie 2 and 3). However, it was the dynamic epicardial expression or migration that preceded the endothelial cell migration into the wound site. The pre-existing mural cell expression moved more slowly alongside the cell body of the endothelial cells, behind the filopodial extension (Supplementary Movie 2 and 3). These imaging data above suggest that two different *pdgfrβ+* populations are present at the wound site during zebrafish heart regeneration: the pre-existing mural cells and a second epicardial cell population. Furthermore, a subcomponent of the epicardium may be dynamically changing gene expression and transforming cell identity.

### Heterogeneous epicardial and mural cells in heart regeneration

We hypothesized that *pdgfrβ+* populations with important roles in zebrafish heart regeneration might have more active gene expression. To further characterize the diverse *pdgfrβ+* cell populations and determine their gene expression signatures, we performed FACS using wildtype adult *Tg*(*pdgfrβ:EGFP*) zebrafish and sorted EGFP positive cells from heart ventricles of uninjured and injured wild type adult zebrafish hearts at 7 dpa to perform scRNAseq. Differentially expressed genes in 15 clusters were identified (Fig. S5A). We found that *pdgfrβ* expression is mainly detected in clusters of epicardial cells/EPDCs/mural cells (cluster 5, 6, 10 and 12) (Fig. S5.1A’). The heterogeneity of *tcf21:nucEGFP+* epicardial cells from uninjured hearts was previously reported (Cao et al., 2016). Therefore, we focused our gene signature analysis on the regenerating hearts. The epicardial/EPDC/mural cell clusters were identified based on the marker gene expressions (312 cells from uninjured hearts and 199 cells from 7 dpa hearts) and isolated and re-clustered into 5 subclusters to determine and analyze the various subpopulations (Fig. S5.1B, C).

When quantifying relative representation of the uninjured vs. injured sample in each cluster, we found the cells from injured heart present more in subcluster 0 (∼45% of injured vs. ∼ 33% of uninjured heart cells) and subcluster 3 (∼10% of injured vs. ∼6% of uninjured heart cells) compared to uninjured heart. On the other hand, uninjured heart cells are present more in subcluster 2 (∼26% uninjured vs. ∼12% of injured heart cells). Subcluster 1 (31% of uninjured vs. ∼28% of injured heart cells) and 4 (∼4% of uninjured vs. ∼6% of injured heart cells) have nearly equal representation from uninjured and injured samples (Fig.5A, A’). By epicardial marker (*tcf21*) and mural cell marker (*rgs5a*) (Cho et al., 2003, Venero Galanternik et al., 2017) gene expressions, subclusters, 0, 1, 2, 3 were found to be epicardial/EPDC subclusters which have very few cells with mural cell marker expression and subcluster 4 was found to be the mural cell subcluster. (Fig. 5B). Interestingly when *pdgfrβ* expression was checked among these subclusters, the mural cell subcluster (subcluster4) showed high *pdgfrβ* expression but there was little to no difference in expression level between mural cells from uninjured and injured hearts. Among epicardial subclusters, cells from injured hearts in subcluster 0 and 3 showed more *pdgfrβ* expression than uninjured heart cells (Fig. 5B). Thus, we selected subcluster 0, which seems most responsive to heart injury and performed Gene Ontology (GO) term analysis based on the differentially expressed genes in this subcluster. After categorization of all enriched GO terms in cluster 0, it was found the functionalities related to tissue development/morphogenesis (25% of all GO terms), peptidase/proteolytic regulation (18%), extracellular matrix (10%), cell movement (9%), wound response/regeneration (5%) are enriched by the differentially expressed genes (Fig. 5C). These functions are well-aligned with the requirements of a regenerating heart where immediately after amputation, extracellular matrix deposition occurs. After that, as the healing process progresses the matrix is dissolved by peptidase/proteolytic activities, and simultaneously cell migration and wound response/regeneration occur.

**Figure 5.**
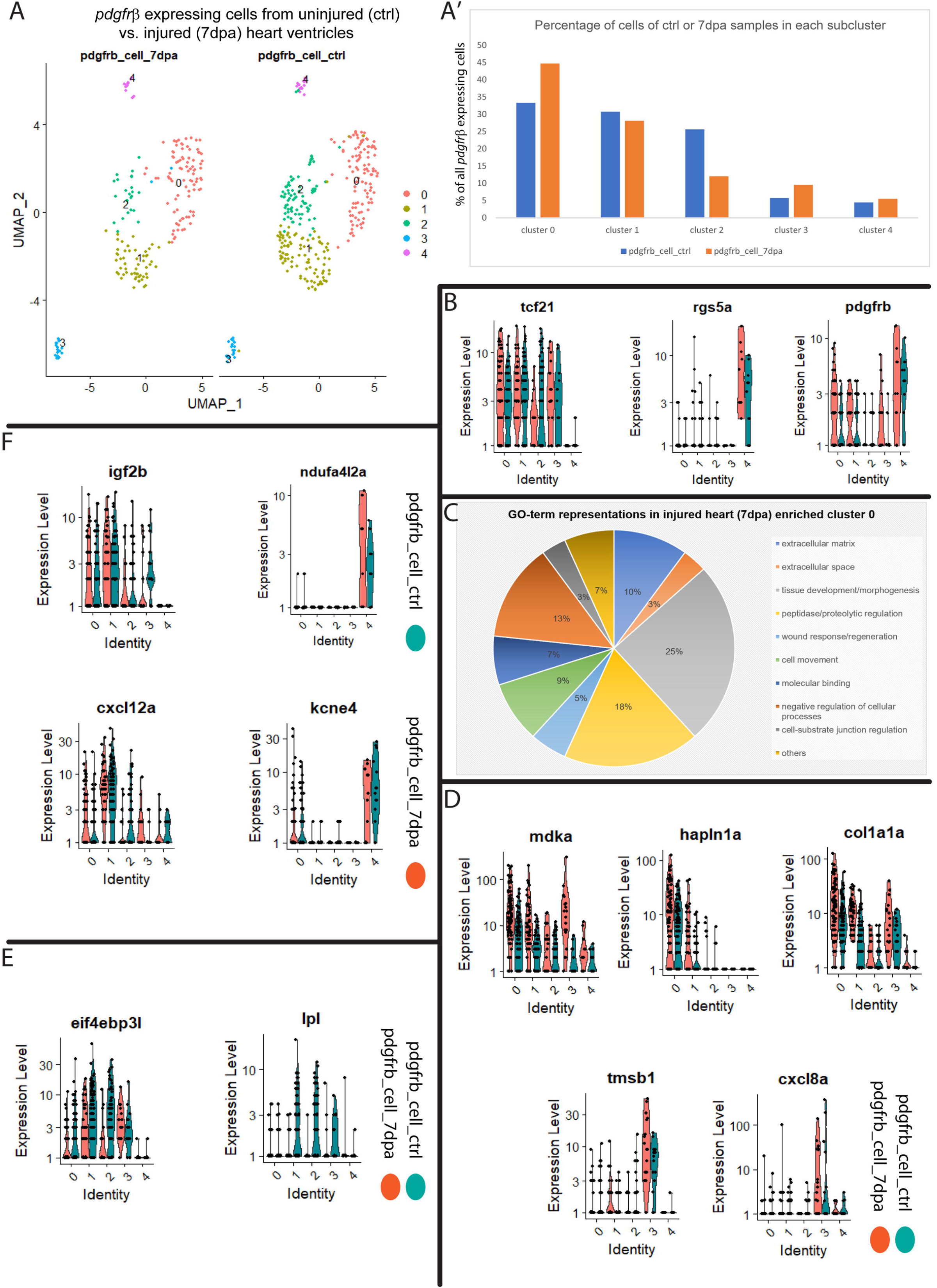
Characterization of the Pdgfrβ+ cells in regenerating hearts after amputation. (A-A’) All cells re-clustered from the FACS sorted *pdgfrβ:EGFP*+ and *pdgfrβ* mRNA expressing epicardial/EPDC and mural cell clusters. The cell are isolated and integrated from 5-7 uninjured and injured (7 dpa) *Tg*(*pdgfrβ:EGFP*; *cxcl12b:*Citrine) fish. (A) The UMAP plot of 5 subclusters of integrated uninjured and 7dpa epicardial/EPDC and mural cells. (A’) Percentages of cells from uninjured control and 7dpa samples in each subcluster, showing relative abundances of cells from each sample in different subclusters. (B) Comparative violin plots (uninjured, blue bars vs. 7dpa, orange bars) for epicardial marker (*tcf21*), mural cell marker (*rgs5a*) and *pdgfrβ* expressions across all subclusters. (C) Categorizations and relative enrichments of Gene Ontology (GO) terms derived from the differentially expressed genes in the subcluster 0. Genes are selected with adjusted p-value < 0.1 and enriched GO-terms were selected with Holm-Bonferroni correction p-value < 0.05. (D) Examples of the differentially expressed genes from subcluster 0 (*mdka, hapln1a, col1a1a*) and subcluster 3 (*tmsb1, cxcl8a*) in respective violin plots. Blue bar indicates uninjured heart and orange bar indicates injured heart (7dpa). (E) Examples of the differentially expressed genes from subcluster 2 (*eif4ebp3l, lpl*). Blue bar indicates uninjured heart and orange bar indicates injured heart (7dpa). (F) Examples of the differentially expressed genes from subcluster 1 (*igf2b, cxcl12a*) and subcluster 4 (*ndufa4l2a, kcne4*) in respective violin plots. Blue bar indicates uninjured heart and orange bar indicates injured heart (7dpa).

Among the genes (from the subcluster 0) which are generally induced in tissue regenerative tissues are *mdka, hapln1a, col1a1a*. Midkine a (*mdka*), a growth factor, was previously shown to be induced during zebrafish heart regeneration (Lien et al., 2006). Hyaleuronan and proteoglycan link protein 1a (*hapln1a*) is predicted to have hyaluronic acid binding activity and was shown to be involved in fin regeneration (Ouyang et al., 2017), epicardial EMT, and heart regeneration (Missinato et al., 2015). Collagen type 1, alpha 1a (*col1a1a*) was also shown to be involved in fin development and regeneration (Duran et al., 2015, Duran et al., 2011, Padhi et al., 2004). Along with subcluster 0, these genes were also seen to be induced in injured heart cells from other subclusters. Thymosin beta 1 (*tmsb1*), Chemokine ligand 8a (*cxcl8a*) are gene candidates from subcluster 3. *tmsb1* is predicted to regulate cell migration, and *cxcl8a* regulates neutrophil migration in response to inflammation due to tissue injury (Powell et al., 2017, Sarris et al., 2012). Translation initiation factor, *eif4ebp3l*, and lipoprotein lipase (*lpl*) are two highly differentially expressed genes in subcluster 2, where more cells from uninjured hearts are present (Fig. 5E). Insulin like growth factor 2b (*igf2b*), chemokine ligand 12a (*cxcl12a*) are differentially expressed genes in the subcluster 1 and the potassium voltage-gated channel, *kcne4*, mitochondrial complex associated protein, *ndufa4l2a* are highly expressed genes in the subcluster 4, where cells from both uninjured and injured hearts are relatively equally included (Fig. 5F). Consistent with our previous findings (Kim et al., 2010), expression of epicardial markers (*tcf21, tbx18, wt1b*), epithelial to mesenchymal (EMT) transition genes (*snai1a, snai1b, snai2, twist1b*) that are induced during heart regeneration and interstitial/extracellular matrix components (*fn1a, fn1b, postnb*) are shown in different violin plots for uninjured vs. injured hearts (Fig. S5.2). Overall, these data suggested that in response to zebrafish heart injury (amputation), a few subpopulations of the heterogeneous *pdgfrβ* expressing epicardial cells (e.g., cells from subcluster 0) more actively express genes for regeneration than pre-existing mural cells (subcluster 4). These results suggest that EPDCs are more actively contributing to heart regeneration than pre-existing mural cells that also migrate into the regenerating area with the endothelial cells.

### *pdgfrβ* is required for heart regeneration

We previously reported that fish treated with Pdgfrβ inhibitor showed defects in revascularization during heart regeneration (Kim et al., 2010). Here, we utilized adult *pdgfrβ* mutants to further examine the requirement of *pdgfrβ* during zebrafish heart regeneration. We observed very few blood vessels in the regenerating area of *pdgfrβ* mutant hearts at 21 and 34 dpa. These vessels were large, had significantly fewer branches, and were mostly devoid of *pdgfrβ+* mural cells. In contrast, sibling-control fish had dense networks of *pdgfrβ+* mural cell-covered coronary vessels in the regenerating region (Fig. 6A). These data suggest that Pdgfrβ signaling plays an essential role during revascularization. The *pdgfrβ* mutants failed to regenerate after ventricular resection and maintained a fibrotic scar (Fig. 6B), confirming that Pdgfrβ signaling is essential for heart regeneration.

**Figure 6.**
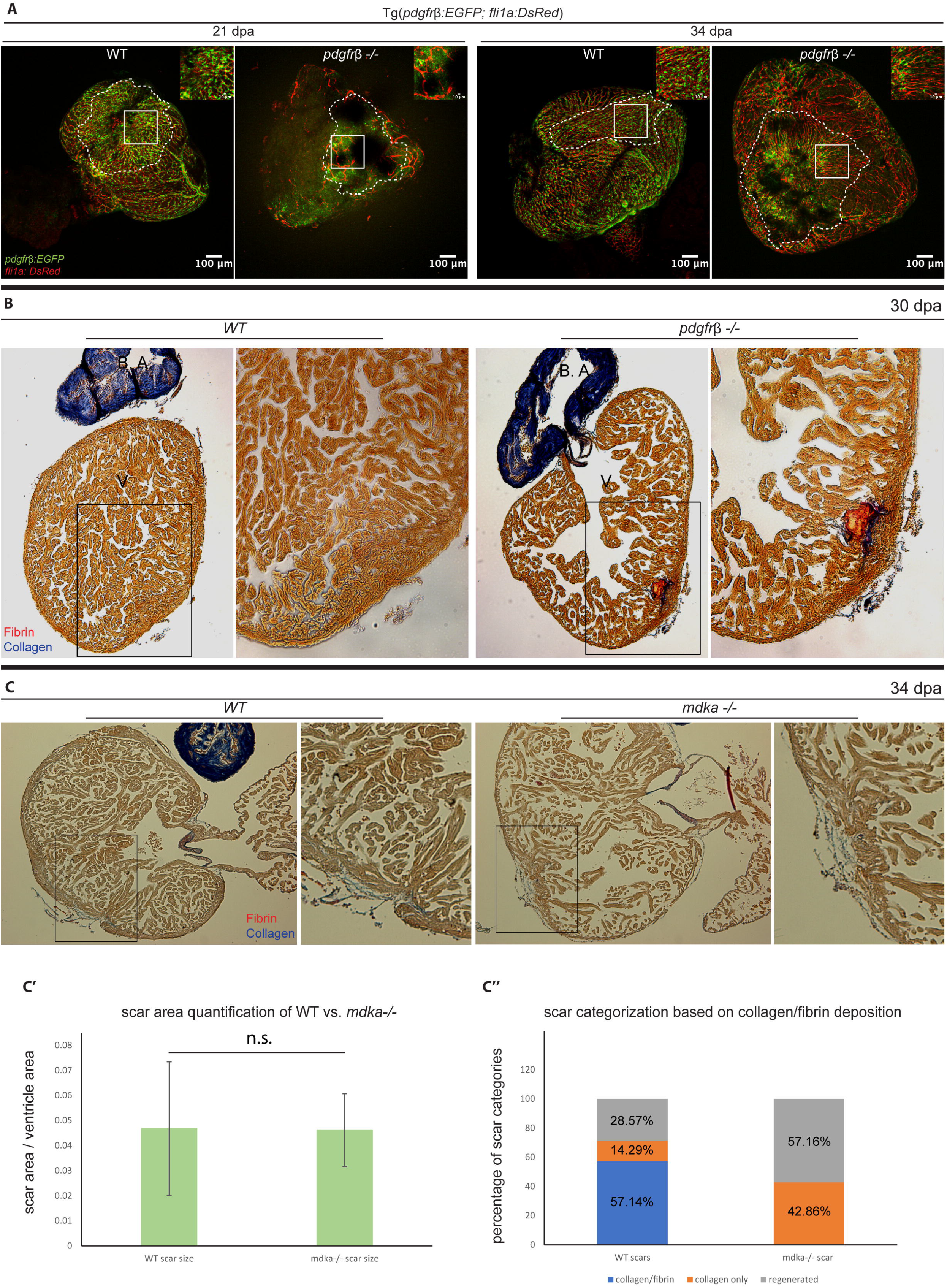
*pdgfrβ* mutant but not *mdka* mutant hearts fail to regenerate and form a fibrotic scar. (A) *pdgfrβ* mutant fish cannot revascularize the regenerating area of the amputated heart. *pdgfrβ*^*+/+*^ fish have dense network of coronary vasculature (*pdgfrβ:EGFP*, green; *fli1a:DsRed*, red) in the regenerating area at 21 and 34 dpa while the *pdgfrβ* mutants have very few coronary vessels with network formation. Unlike WT, the coronary vessels in the *pdgfrβ* mutant lack the *pdgfrβ+* mural cell coverage in regenerating area (n = 4). (B) AFOG staining of heart sections of WT and *pdgfrβ* mutants at 30 dpa. n=7. (C) AFOG staining of WT and *mdka* mutant heart at 34 dpa. n=7. fibrin, red; collagen, blue. (C’) Quantification of scar area in WT vs mdka mutants. n.s. not significant. (C”) Quantification of scar categorization. Percentage of collagen/fibrin deposition is shown in blue, collagen only in orange, and regenerated in grey.

### *midkine-a* (*mdka*) mutants do not show significant heart regeneration defects

*mdka* was found in subcluster 0 and showed increased expression in 7 dpa hearts. We confirmed the upregulation of *mdka* in EDPCs by qRT-PCR using FACS sorted *pdgfrβ*:*EGFP+* cells and absence of *mdka* in *mdka* mutants (Nagashima et al., 2020) by *in situ* hybridization and qRT-PCR (Fig. S6A, B, C). Midkine (mdk) has been shown to play important roles in tissue regeneration in many different organs (Ang et al., 2020, Ikutomo et al., 2014, Nagashima et al., 2020, Tsai et al., 2020). However, in contrast to *pdgfrβ* mutant hearts, *mdka* mutants did not show significant defects in heart regeneration (Fig. 6C) and collagen deposition (Fig. 6C’, C”) despite its strong expression.

## DISCUSSION

The cell compositions, functions and origins of the mural cells in the hearts still remain incompletely understood. Focusing on *pdgfrβ*, a well-known pericyte marker, we determined the heterogeneity of cardiac mural cells and epicardial derived cells (EPDCs) in zebrafish heart during development and regeneration and how Pdgfrβ signaling affects these different cell populations. During heart development, *pdgfrβ* and *cxcl12b* double positive (*pdgfrβ*+;*cxcl12b*+) cells line coronary arteries while *pdgfrβ*+ only cells surround non-arterial vessels. In adult regenerating hearts, *pdgfrβ+* cells are identified in pre-existing mural cells and EPDCs. These *pdgfrβ+* cells behave differently during heart development and regeneration. The use of scRNAseq and confocal and live imaging allowed us to delineate unique gene expression signatures in different populations, revealing their different functions.

*pdgfrβ*+ mural cells enveloped coronary arteries also expressed *cxcl12b*. Although very few *pdgfrβ*+;*cxcl12b*+ double positive cells were captured in our analysis, smooth muscle gene expression in these double positive cells suggests that they are more differentiated than the *pdgfrβ*+ only cells. Interestingly, the association of these *pdgfrβ*+;*cxcl12b*+ mural cells with coronary arteries does not depend on Pdgfrβ signaling while all non-arterial vessels lose *pdgfrβ*+ mural cells in *pdgfrβ* mutants. Recent scRNAseq data of mouse brain revealed two subclasses of mural cells; pericytes are in a continuum with venous smooth muscle cells which are distinct from arteriole or arterial smooth muscle cells (Vanlandewijck et al., 2018). It is possible that *pdgfrβ*+;*cxcl12b*+ mural cells are similar to arteriole or arterial smooth muscle cells which are different from the *pdgfrβ*+ only cells surrounding the wide large vessels that are likely the coronary veins. In mice, smooth muscle cells along the coronary arteries are derived from *pdgfrβ+* pericyte progenitors after the blood flow starts while the smooth muscle cells around the coronary arteries fail to form in Pdgfrβ knock out mice (Volz et al., 2015). However, these *pdgfrβ*+;*cxcl12b*+ double positive cells remained associated with coronary arteries in *pdgfrβ* mutant fish, suggesting a novel mechanism exists in maintaining the mural cell association with coronary arteries. One likely mechanism is that Cxcl12-Cxcr4 signaling which has been shown to regulate the preferential recruitment of smooth muscle cells to arteries (dorsal aorta in fish) (Stratman et al., 2020). Our scRNAseq also identified several signaling pathways that could mediate this Pdgfrβ independent mural cell association with coronary endothelial cells and this will be pursued in future investigation.

The functions of Pdgfrβ signaling and *pdgfrβ*+ cells have been implicated to play important roles in the development and regeneration of different tissues and organs. Consistent with our finding, intra-myocardial delivery of PDGF-BB provides myocardial protection and improve ventricular functions (Hsieh et al., 2006a, Hsieh et al., 2006b). Recently it was demonstrated that extracellular matrix (ECM) derived from *pdgfrβ*+ myoseptal and perivascular cells prevents scarring and promote axon regeneration of zebrafish spinal cord in a Pdgfrβ signaling dependent manner (Tsata et al., 2021). Consistent with this finding, our scRNAseq analyses identified significant changes (10% of all GO-term enrichment) in genes encoding ECM components. How Pdgfrβ signaling regulates the ECMs in the regenerating heart and shape the regenerative environment will be of interest for future studies.

*mdka* was identified by our scRNAseq and previously by microarray gene expression profiling (Lien et al., 2006) as an upregulated gene in regenerating hearts. *mdka* was shown to play an important role in neural regeneration in retina by regulating cell cycle progression of Müller glia (Nagashima et al., 2020), epimorphic regeneration (Ang et al., 2020), and regeneration after skeletal muscle injury (Ikutomo et al., 2014). Furthermore, it was recently reported that Midkine (Mdk) could regulate wound epidermis development and inflammation during the initiation of limb regeneration (Tsai et al., 2020). *mdka* is highly expressed in epicardium and EDPCs after amputation and this led us to examine its role during heart regeneration. We did not observe any significant defects in heart regeneration and increased fibrotic scar remains at the injury site at 34 dpa compared to controls. However, we cannot exclude the possibility that Mdka and Mdkb have redundant functions and *mdkb* can compensate the loss of *mdka*. Examining the phenotypes of double mutants is of interest for future investigation; however, is beyond the scope the current manuscript.

## MATERIALS and METHODS

### Fish lines

The following zebrafish lines were raised and maintained at Children’s Hospital Los Angeles (CHLA) under standard conditions of care and with CHLA IACUC oversight. IACUC approved all experimental procedures used in this study. The fish lines have been published in (Villefranc et al., 2007, Lawson and Weinstein, 2002), (Ando et al., 2016), (Vanhollebeke et al., 2015), (Kok et al., 2015), (Nagashima et al., 2020), (Bussmann et al., 2011), (Kikuchi et al., 2011), (Mosimann et al., 2011).

| Fish Line | Reference |
| --- | --- |
| *Tg(fli1a.ep:DsRedEX)<sup>um13</sup>* also known as *fli1a:DsRed* | (Villefranc et al., 2007) |
| *Tg(fli1a:EGFP)* | (Lawson and Weinstein, 2002) |
| *TgBAC(pdgfrβ:EGFP)* | (Ando et al., 2016) |
| *TgBAC(pdgfrβ:Citrine)* | (Vanhollebeke et al., 2015) |
| *pdgfrβ<sup>um148</sup>* | (Kok et al., 2015) |
| *Mdka<sup>mi5001</sup>* | (Nagashima et al. (2020) |
| *Tg(cxcl12b:Citrine)* | (Bussmann et al., 2011) |
| *TgBAC(tcf21:Cre-ERT2)<sup>pd42</sup>* also known as<br>*tcf21:CreERT2* | (Kikuchi et al., 2011) |
| *Tg(-3.5ubi:loxP-EGFP-loxP-mCherry)* also known as<br>*ubi:Switch* | (Mosimann et al., 2011) |

Lineage tracing *tcf21:CreERT2* lineage tracing was performed as described in (Harrison et al., 2015) with *pdgfrβ:Citrine* as the mural cell marker. *Tg(pdgfrβ:Citrine; tcf21:CreERT2; ubi:loxp-EGFP-loxp-mCherry)* fish were generated and the embryonic epicardial cells were labeled by mCherry by activating *tcf21:CreERT2* by administering 4-hydroxytamoxiphen (4 OHT) during first 5 days of the embryonic development. 13μg/ml 4 OHT was added in E3 medium and the medium was changed daily. After the treatment the fish were raised into adulthood. The adult hearts carried all the progenies derived from the embryonic labeled epicardium. *pdgfrβ:Citrine* was used to identify the mural cells. The percentage of mCherry labeled, Citrine positive cells were quantified.

### Angiography Dye Injection and Amputation

Dextran cascade blue dye was injected through the retro-orbital sinus as described (Harrison et al., 2015). Briefly, injection was done 2.5-3.0 hours before heart extraction. Zebrafish were anesthetized in a tricaine solution and placed horizontally on a sponge so that its right eye was facing up. A Hamilton syringe loaded with 4pL Dextran Cascade Blue was inserted at 30° to the plane of the sponge, about 2mm deep at the 7 o’clock position of the right eye socket (Pugach et al., 2009). The amputation experiments were performed as described in Poss et al., 2002. The adult zebrafish was anaesthetized by administering 0.168 mg/ml Tricaine in the system water in beaker. The anaesthetized fish was placed in the dissection grove and using small scissor and forceps opened the chest cavity. The apical portion of the heart ventricle (∼ 20% of the ventricle) was amputated. The fish were then kept in shallow system water for a while until blood flow stops from the operated site and it recovers regular movement.

### Confocal Imaging

Hearts were immobilized in 1% low melting point agarose in PBS. The heart was oriented with the ventricle at the center of the image, the bulbus arteriosus at the top, and the atrium on the left. Z-stack images were collected utilizing a Zeiss 710 confocal microscope. Z-stack images were then converted to a maximum intensity projection before further processing/quantification. ImageJ/ Fiji software was utilized for quantification.

### Live Imaging

Hearts from transgenic zebrafish *Tg(pdgfrb:EGFP; fli1a:DsRed)* were amputated and allowed in recover *in vivo* for 5-10 days after which hearts were removed into imaging media [L15 (300ml), 10% FCS/FBS, 100ug/ml Primocin, 1.25mM CaCl2, 800mg/L glucose, Pen/Step]. Hearts were cleaned of any external blood or attached tissue debris. The imaging microfluidic device was prepared as described [(Yip et al., 2019); Harrison *et al*. in preparation] and four hearts were then placed into the imaging wells under ringer’s solution. The device was sealed and mounted onto a Zeiss cell observer system equipped with a Hamamatsu ORCA-flash4.0LT and Colibri 7 LED light source for live imaging. The imaging device was maintained at a temperature of 28.5°C with a PeCon atmospheric control stage and cover. Imaging media was perfused through the imaging wells at a rate of 0.5 mL/hr and imaging was carried out for 72-120 hours. Acquisitions were carried out to capture a z stack of 35-50 images 6um part every 30 minutes. Collected images were cropped (in z and time) and deconvolved (using AutoQuant). ImageJ/Fiji was used to produce the final moves with the aid of a Gaussian-based focusing macro and a z sub-selection macro (available on request).

### Quantification

#### Mural cell association

Mural cell association with the coronary vessels were quantified as the mural cell number/unit vessel length. Using the ImageJ software, freehand lines were drawn along the length of the randomly selected vessels of certain size (e.g., Fig. 2B’) or from certain regions of the heart ventricles (e.g., Fig. 1B). The length (by pixel numbers) of the freehand line was measured by ImageJ and the number of *pdgfrβ+* cells along the line were counted manually. Then the *pdgfrβ+* cell numbers were divided by the freehand line’s length to get the measurement of the mural cell number/ unit vessel length. Paired t-test was used to quantify the significance of the value differences between different samples.

Ventricle coverage by coronary vessels: Using ImageJ freehand selection tools, areas covered by coronary vessels following the tips of the vessels were selected in heart ventricles. Respective areas were measured and the areas following the coronary vessels were divided by whole ventricle area and corresponding percentages were quantified (Fig. 2A’). One-way ANOVA was used to quantify the significance of the value differences between *pdgfrβ*^*+/+*^, *pdgfrβ*^*+/-*^ and *pdgfrβ*^*-/* -^hearts.

#### *pdgfrβ* expression during regeneration

*pdgfrβ* expression was quantified by measuring the area percentage of the *pdgfrβ* expressing region against the area of whole apical view of the ventricle (Fig. 3A’). The area of the *pdgfrβ* expressing region (marked by the white small-dotted line in Fig. 3A) was quantified by using freehand selection tool of the Image J software. One-way ANOVA was used to quantify the significance of the measurement difference between 3dpa and 7dpa.

### Heart tissue dissociation into single cell suspension

The heart ventricle was immersed in the modified Tyrode’s solution (Tessadori et al., 2012) on ice. After washing blood from the tissue, the ventricle was torn open/into small pieces by forceps and transferred in to Ca^2+^ free modified Tyrode’s solution (Tessadori et al., 2012) at 30° C. The tissue was then washed 2X times with ice-cold Ca^2^+ free modified Tyrode’s solution. After this the tissue was treated with the digestion mix (500µl/ 5 hearts) containing Liberase (500 CDU); Elastase (3.1 U) and DNasel (32 U) at 33°-35° C for ∼ 15 minutes with continuous stirring. Next the digestion was stopped by adding ice-cold Ca^2^+ free modified Tyrode’s solution containing 10% Fetal Bovine serum and DNase1 (32 U). The solution was then filtered through 40-micron sieve and the cells are pelleted by centrifuging at 500Xg for 5 minutes at 4° C. The cells were washed with ice-cold Ca^2+^ free modified Tyrode’s solution containing 30% Fetal Bovine serum. The cells were then precipitated again and redissolved in appropriate volume of ice-cold Ca^2+^ free modified Tyrode’s solution containing 30% Fetal Bovine serum to maintain proper cell density for single cell cDNA library preparation or qRT-PCR.

### Fluorescence Activated Cell Sorting (FACS) sorting of *pdgfrβ:EGFP* cells

*pdgfrβ:EGFP* cells were FACS sorted from *Tg*(*pdgfrβ:EGFP*; *cxcl12b:Citrine*) 6 month old wild type and *pdgfrβ*^*-/-*^ fish. *pdgfrβ:EGFP* cells were FACS sorted from ∼10, *Tg*(*pdgfrβ:EGFP*; *cxcl12b:Citrine*) 18 months old uninjured or injured (7 dpa) fish. Heart ventricles were dissociated into single cell suspension following abovementioned method for each sample. *pdgfrβ:EGFP* cells were FACS sorted using BD FACSAria cell sorter. The live cells were sorted by filtering out dead cells with DAPI staining. EGFP only, DsRed only transgenic samples were used as positive control and wildtype fish without any transgenic reporter were used as the negative control. During sorting the FACS gates were set to maximize inclusion of all GFP positive cells (low to high EGFP signals) to isolate all possible EGFP expressing cell populations. Although this wide gate set-up leads to inclusion of other cell types (e.g., cardiomyocytes) which do not express *pdgfrβ:EGFP*, but possibly have background fluorescence. The cells were collected in a Ca^2+^ free modified Tyrode’s solution containing 30% Fetal Bovine serum and kept on ice. It was centrifuged at 500Xg for 5 minutes at 4° C to concentrate the sample for downstream procedures (e.g., Single cell RNA sequencing, qRT-PCR).

### Single cell RNA sequencing

For the generation of single-cell gel beads in emulsion, cells were loaded on a Chromium single cell instrument (10x Genomics) per manufacturer’s protocol. In brief, single-cell suspension of cells in 0.4% BSA-PBS were added to each channel on the 10x chip. Cells were partitioned with Gel Beads into emulsion in the Chromium instrument where cell lysis and barcoded reverse transcription of RNA occurred following amplification. Single cell RNA-Seq libraries were prepared by using the Chromium single cell 3’ library and gel bead kit v3 (10x Genomics). Sequencing was performed on HiSeq platform (Illumina), and the digital expression matrix was generated using the Cell Ranger pipeline (10x Genomics). 1833 (99,466 mean reads per cells) and 1504 *pdgfrβ:EGFP* cells (67,463 mean reads per cell) were sequenced for control and *pdgfrβ* mutant hearts. After removing low quality cells, we analyzed 955 and 1126 cells from control and *pdgfrβ* mutant hearts respectively. From combined dataset of control and *pdgfrβ* mutant hearts (955+1126 cells), 274 *pdgfrβ* only and 126 *pdgfrβ; cxcl12b* cells were identified and analyzed. 5826 cells from uninjured hearts and 3127 cells from regenerating WT adult hearts with an average of 29,327 and 66,193 reads per cell, respectively. After removing low quality cells, finally we analyzed 1,631 cells from uninjured heart and 1097 cells 7 dpa hearts. From combined dataset of uninjured and injured hearts (1631+1097 cells), 511 *pdgfrβ* cells or epicardial/EPDC/mural cells were identified and analyzed by Seurat R package.

### Single cell RNA sequencing data analysis

To identify different cell types and find signature genes for each cell type, the R package Seurat (version 3.2.3) was used to analyze the digital expression matrix. Cells with less than 100 and greater than 2500 unique feature count and greater than 25% mitochondrial expression were removed from further analysis. Seurat function NormalizeData was used to normalize the raw counts. Variable genes were identified using the FindVariableGenes function. The Seurat ScaleData function was used to scale and center expression values in the dataset for dimensional reduction. Principal component analysis (PCA), t-distributed stochastic neighbor embedding (t-SNE), and uniform manifold approximation and projection (UMAP) were used to reduce the dimensions of the data, and the first 2 dimensions were used in plots. The FindClusters function was later used to cluster the cells. The FindAllMarkers function was used to determine the marker genes for each cluster, which were then used to define cell types. Also, known cell type marker expressions were determined across different clusters to assign the cell type to a cluster. We integrated the wildtype and *pdgfrβ*^*-/-*^ dataset together. Also, we integrated uninjured and injured (7dpa) datasets by using pairwise anchors by Seurat. After clustering cells based on the differentially expressed genes, cell types were identified by marker genes expressions. For Gene Ontology (GO) term analysis, differentially expressed genes in a cluster were selected with adjusted p-value < 0.1. The selected genes were then put in a list form at http://www.zebrafishmine.org website for GO term analysis. Enriched GO-terms were generally selected after running Holm-Bonferroni or Benjamini Hochberg corrections with p-value <0.05. The genes from our dataset under each GO-term were identified and looked for selected genes’ expression in individual cells of the samples (e.g., wildtype vs. *pdgfrβ*^*-/-*^) by heatmap (Fig. S3.1, C’) to validate the GO-term enrichment. Different plots (e.g., violin plots, ridge plots, feature plots) were made following available default Seurat codes. In the ridge plots, for individual genes, number of cells expressing the gene at certain expression level (expression level > 0 and expression level > 1) was determined and the percentages were calculated against total cell number of the respective clusters (e.g., *pdgfrβ*+ only cluster or *pdgfrβ+*; *cxcl12b+* cell cluster) for wildtype or *pdgfrβ* mutant conditions.

### Quantitative RT-PCR (qRT-PCR)

FACS sorted *pdgfrβ:EGFP* cells or *fli1a:DsRed* cells or whole ventricle samples were collected in Trizol for RNA extraction by Trizol-chloroform method. The RNAs were precipitated overnight at -20° C in isopropanol and washed with 70% alcohol and dissolved in DEPC treated water. cDNAs were made from the RNA by using the SuperScript® III First-Strand Synthesis System. The qRT-PCR were performed using Applied Biosystems® SYBR® Green PCR Master Mix. Gene expression fold change are normalized with the housekeeping gene *rpl13*. The mean cycle threshold (Ct) values of each sample triplicates were calculated and 2^-Ct^ values were calculated for each sample to determine expression fold changes.

qRT PCR primers:

*rpl13a* forward primer 5’ GGTGTGAGGGTATCAACATCTC 3’

*rpl13a* reverse primer 5’ GTGATATGATCCACGGGAAGG 3’

*mdka* forward primer 5’ GGGAACTAAAGGGAGCAAGAG 3’

*mdka* reverse primer 5’ CTGAACAACACAGAGTGGAGAT 3’

*mdkb* forward primer 5’ TCCCATGCAACTGGAAGAAG 3’

*mdkb* reverse primer 5’ CCTGTAGTAGTGTCACATTCGG 3’

*pdgfrβ* forward primer 5’ TATGTGTGCACCGAGAAGAAG 3’

*pdgfrβ* reverse primer 5’ GAACCACACGTCAGGATCAG 3’

## ACKNOWLEDGEMENTS

We thank Drs. Koji Ando and Christer Betsholtz for discussions and communications prior to the submission, and Dr. Didier Stainier for the *pdgfrβ:Citrine* line and Drs. Koji Ando, Hiro Nakajima and Naoki Mochizuki for the *pdgfrβ:EGFP* line And Dr. Ken Poss for the *tcf21:dsRed* line. We also thank the TSRI Cellular Imaging, FACS and Single-Cells, Sequencing and CyTOF (SC2) cores.

