## Supplemental Figures for "Heterogeneous *pdgfrβ+* cells regulate coronary vessel development and revascularization during heart regeneration"

**Supplemental information**

**Supplementary Figure 1. *pdgfrβ* expression in the developing zebrafish heart.**

(A) Representative images of *pdgfrβ:Citrine; fli1a:dsRed* hearts used for quantification during coronary vessel development. Proximal (near) and distal (away) to atrioventriculur canal (AVC) areas are highlighted with boxes at 55, 60, 69 and 76 dpf. Arrowheads indicate the *pdgfrβ*+ mural cells. Zoomed in images are on the right. (B-B'') *pdgfrβ* expression in early juvenile hearts (33 dpf). Double (B) and single (B', B'') channel images of *pdgfrβ:Citrine* and *tcf21:DsRed.* A, atrium, V, ventricle, AVC, white arrow. In bulbus arteriosus (B.A.), *tcf21* signal is weak and does not overlap with the *pdgfrβ.* n = 4. (C) Quantification of *pdgfrβ*+ cells derived from *tcf21* lineage-traced cells. In *TgBAC(pdgfrβ:Citrine); tcf21:CreERt2; ubi:loxp-EGFP-loxp-mCherry* fish, ~76.15% of the *pdgfrβ*+ mural cells were co-labelled with *tcf21* lineage-traced cells at 87 dpf. ~23.85% of the *pdgfrβ*+ mural cells were not *tcf21* lineage traced. Error bars, standard deviation of mean. T- test (***p < 0.001), n =3.

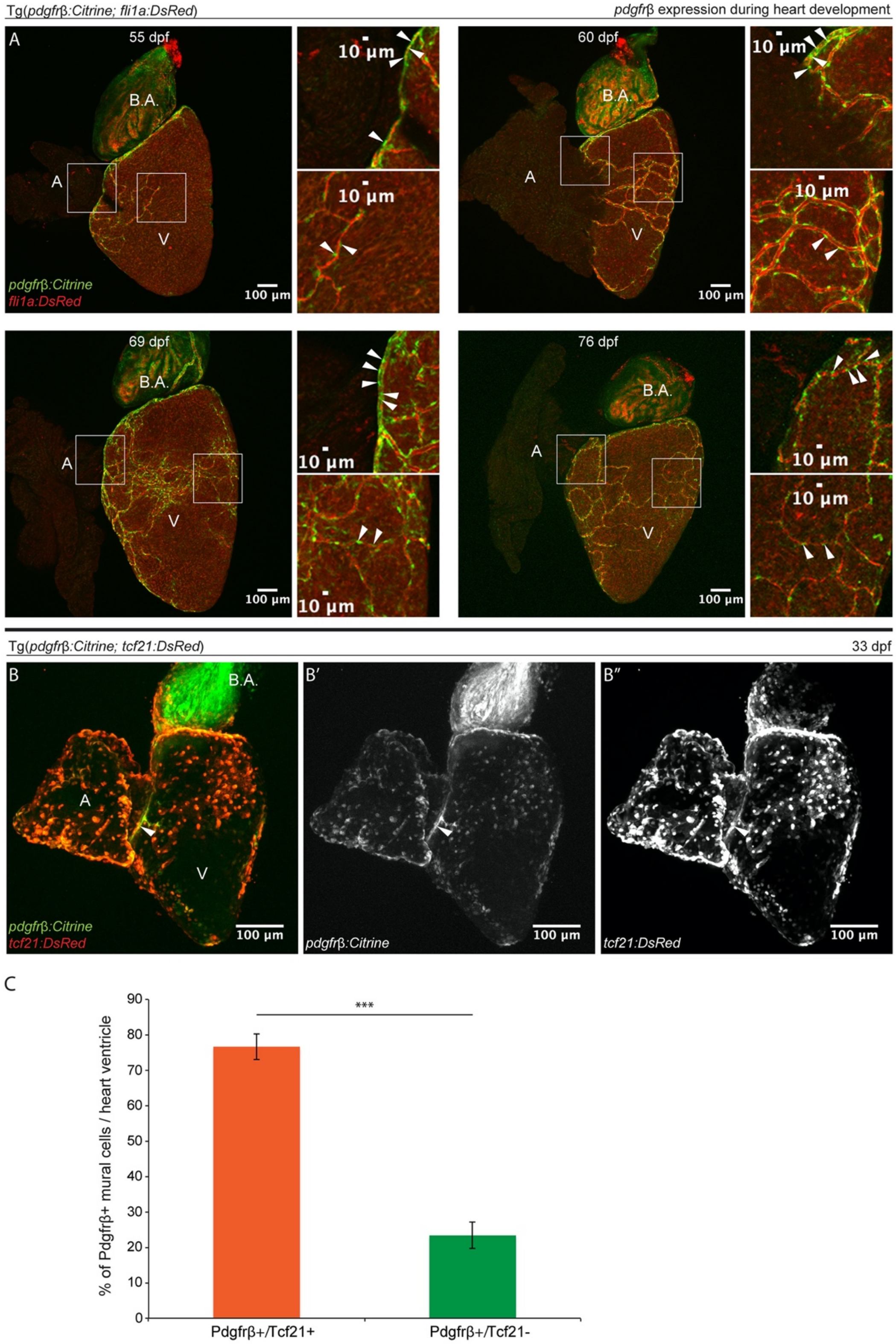

Tg(pdgfrβ:Citrine; fli1a:DsRed)
pdgfrβ expression during heart development
A
55 dpf
60 dpf
69 dpf
76 dpf
B.A.
A
V
pdgfrβ:Citrine
fli1a:DsRed
100 µm
10 µm
Tg(pdgfrβ:Citrine; tcf21:DsRed)
33 dpf
B
B′
B″
pdgfrβ:Citrine
tcf21:DsRed
C
% of Pdgfrβ+ mural cells / heart ventricle
0
10
20
30
40
50
60
70
80
90
***
Pdgfrβ+/Tcf21+
Pdgfrβ+/Tcf21-

**Supplementary Figure 2. Pdgfrβ regulates mural cell number, association, and development of the blood vessels in the brain and coronary vessels.**
(A) *pdgfrβ* loss-of-function mutation affects mural cell recruitment to the brain blood vessels. In adult *pdgfrβ* mutant (*pdgfrβ$^{-/-}$*) fish (224dpf), the brain has significantly decreased mural cell *[Tg(pdgfrβ:EGFP)]* on the blood vessels *[Tg(fli1a:DsRed)].* There is a decline in the vessel density, and the vessels become dilated (indicated by the white arrow and the line showing vessel diameter). (B) In *pdgfrβ* mutant fish *(pdgfrβ$^{-/-}$),* scattered isolated endothelial cells *[Tg(fli1a:GFP)]* are found which fails to form continuous coronary vessels at different developmental stages (98 dpf, 167 dpf). (C-C') In *Tg(pdgfrβ:EGFP; fli1a:DsRed)* fish, the large coronary vessels *[Tg(fli1a:DsRed)]* are classified based on their diameter and appearance into the wide large vessels (yellow arrow) and the narrow large vessels (white arrow). The wide large vessels appear to have less mural cell *[Tg(pdgfrβ:EGFP)]* association than narrow large vessels. (C) In *pdgfrβ* mutant fish *(pdgfrβ$^{-/-}$)* with moderate coronary vessel development, both types of large vessels are formed. While the narrow large vessels (white arrow) maintain the mural cell *[Tg(pdgfrβ:EGFP)]* association, the mural cells around the wide large vessels (yellow arrow) decrease significantly (C'). (D). Quantification of the diameter of large and small coronary vessels. Large coronary vessels (both wide and narrow large vessels) diameter becomes smaller in *pdgfrβ* mutant ventricles. While the mean diameter of the wide large vessels decreases from ~27.6 mm in the *pdgfrβ$^{+/+}$* fish to ~12.9 μm in *pdgfrβ$^{-/-}$* fish, the narrow large vessels become thin from ~13.3 μm diameter to ~8.3 μm (n = 3 hearts, 3 X 3 quantifications from each heart) T-test (***p <0.001, **p < 0.01).

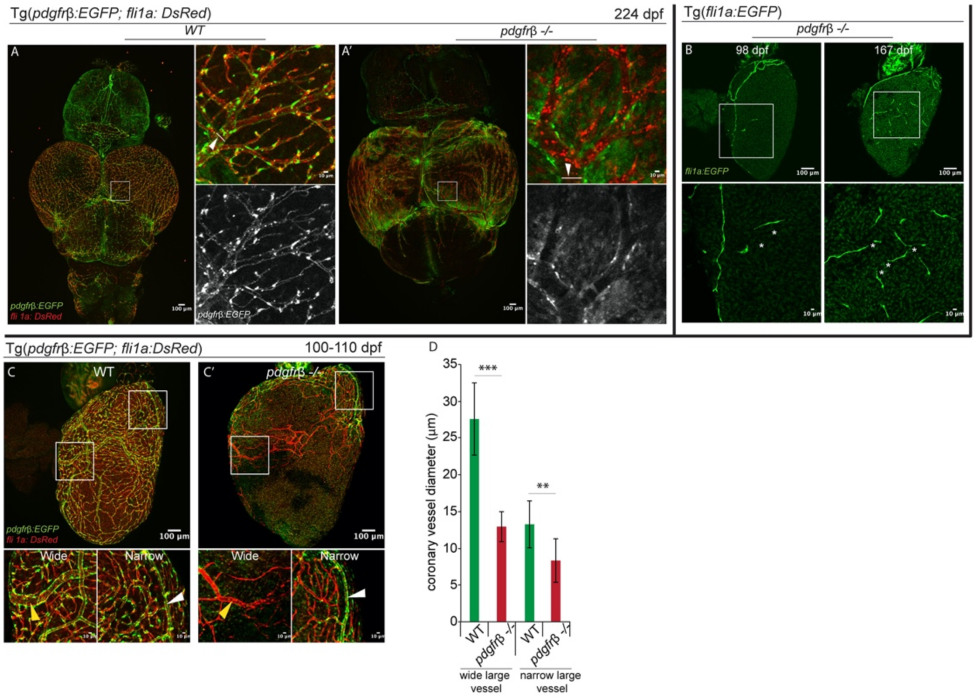
Tg(pdgfrβ:EGFP; fli1a: DsRed)
224 dpf
WT
pdgfrβ -/-
A
A'
pdgfrβ:EGFP
fli 1a: DsRed
100 µm
10 µm
Tg(fli1a:EGFP)
pdgfrβ -/-
B
98 dpf
167 dpf
fli1a:EGFP
Tg(pdgfrβ:EGFP; fli1a:DsRed)
100-110 dpf
C
WT
C'
pdgfrβ -/-
Wide
Narrow
D
coronary vessel diameter (µm)
35
30
25
20
15
10
5
0
***
**
WT
pdgfrβ -/-
wide large vessel
narrow large vessel

**Supplementary Figure 3.1. Identification and characterization of the epicardial cells** (A-A') *pdgfrβ:EGFP* cells are FACS isolated from *Tg*(*pdgfrβ:EGFP*; *cxcl12b:Citrine*) fish. (A) UMAP plot of all isolated EGFP+ cells (from the wild type and *pdgfrβ*$^{-/-}$ sample integrated together), which form 9 clusters. (A') Feature plots of *pdgfrβ* and *cxcl12b* genes that show the cells in the cluster 3 and 6 express *pdgfrβ*. Only cluster 6 cells express *cxcl12b*. (B-B') Cell type identification by marker gene expression. (B) Violin plots are showing EGFP mRNA from the *pdgfrβ:EGFP* reporter is mostly expressed in the cluster 3 and 6 which also specifically express epicardial marker genes (*tcf21*, *tbx18*, *wt1a*). Cluster 0, 1, 2, 4, 5 and 7 highly express cardiomyocyte marker *cmlc1*. Cluster 8 highly express red blood cell marker *hbba1*. (B') UMAP of all the isolated cells showing the cell clusters according to cell types. (C-C') Characterization of the cluster 3 cells (the *pdgfrβ*+ only cell cluster). (C) Cluster 3 cells are isolated and re-clustered in to 5 clusters. Percentages of the wild type and *pdgfrβ*$^{-/-}$ cells in each cluster are plotted in the bar chart. Cluster 0, 2 are over-represented by *pdgfrβ*$^{-/-}$ cells and cluster 1, 3 are over-represented by the wildtype cells. Cluster 4 has similar representations from both samples. (C') Gene Ontology (GO) term enrichment analysis of all the clusters. Three representative GO-terms are selected for each cluster and expression of 1-2 candidate genes for each GO-term is shown in all the *pdgfrβ*+ only cells by heatmap. The plot is arranged according to wild type vs. *pdgfrβ*$^{-/-}$ conditions. For GO-term analysis differentially expressed genes for each cluster have been selected using adjusted p-value cut-off <0.1 and then the GO-terms were filtered using Benjamini Hochberg correction (p value < 0.05).

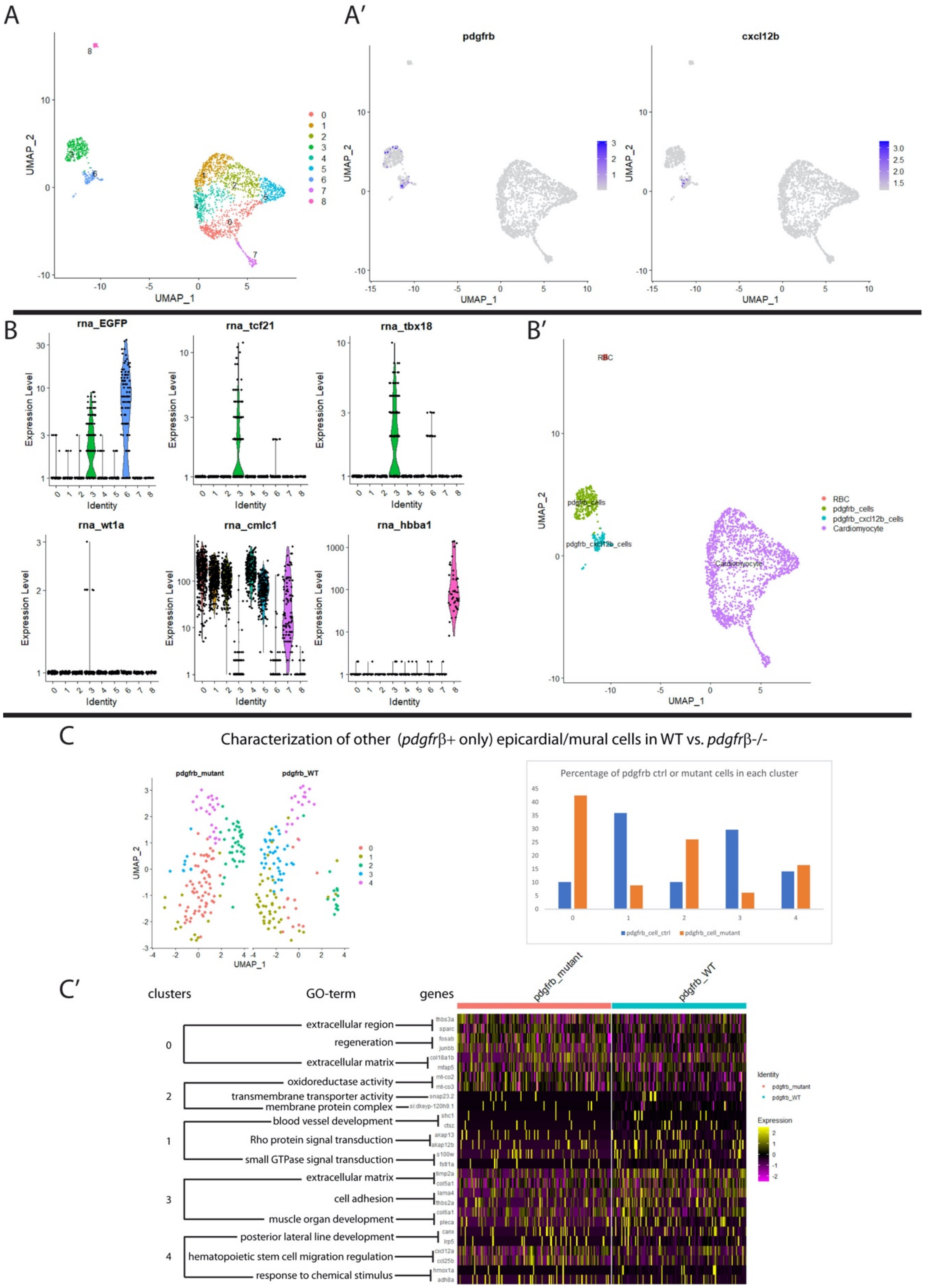
A
A′
pdgfrb
cxcl12b
UMAP_1
UMAP_2
B
rna_EGFP
rna_tcf21
rna_tbx18
rna_wt1a
rna_cmlc1
rna_hbba1
Expression Level
Identity
B′
RBC
pdgfrb_cells
pdgfrb_cxcl12b_cells
Cardiomyocyte
C
Characterization of other (pdgfrβ+ only) epicardial/mural cells in WT vs. pdgfrβ-/-
pdgfrb_mutant
pdgfrb_WT
Percentage of pdgfrb ctrl or mutant cells in each cluster
pdgfrb_cell_ctrl
pdgfrb_cell_mutant
C′
clusters
GO-term
genes
extracellular region
regeneration
extracellular matrix
oxidoreductase activity
transmembrane transporter activity
membrane protein complex
blood vessel development
Rho protein signal transduction
small GTPase signal transduction
extracellular matrix
cell adhesion
muscle organ development
posterior lateral line development
hematopoietic stem cell migration regulation
response to chemical stimulus
Identity
Expression

**Supplementary Figure 3.2. The ridge-plot of different marker gene expression in the *cxcl12b*+;*pdgfrβ*+ cell cluster or the *pdgfrβ*+ only cell cluster of wild type vs. *pdgfrβ*$^{-/-}$ mutant fish.** (A-A') The ridge plots for the smooth muscle marker genes (*acta2*, *myh11a*, *tagln*) and smooth muscle cell differentiation regulator *notch3* in *pdgfrβ*+ only cell cluster. (B) The epicardial cell markers (*tcf21*, *tbx18*) are expressed more in the *pdgfrβ*+ only cell cluster with little expression in the *cxcl12b*+, *pdgfrβ*+ cluster cells. *pdgfrβ*$^{-/-}$ cells have less expression of these markers in the *cxcl12b*+; *pdgfrβ*+ cluster. In the *pdgfrβ*+ only cell cluster, *pdgfrβ*$^{-/-}$ cells express *tbx18* in proportionately more cells at similar level as the wildtype cells. *tcf21* expression decreases in the *pdgfrβ*$^{-/-}$ cells. *wt1a* rarely expresses in both of these clusters and there is not much difference between the wildtype cells and *pdgfrβ*$^{-/-}$ cells. (C) Other mural cell markers (*rgs5a*, *cd248a*, *kcne4*). The mural cell marker *kcne4*, broadly express in both *cxcl12b*+, *pdgfrβ*+ cluster and *pdgfrβ*+ only cluster. All these markers expressed less in the *pdgfrβ*$^{-/-}$ cells than in wild type cells.

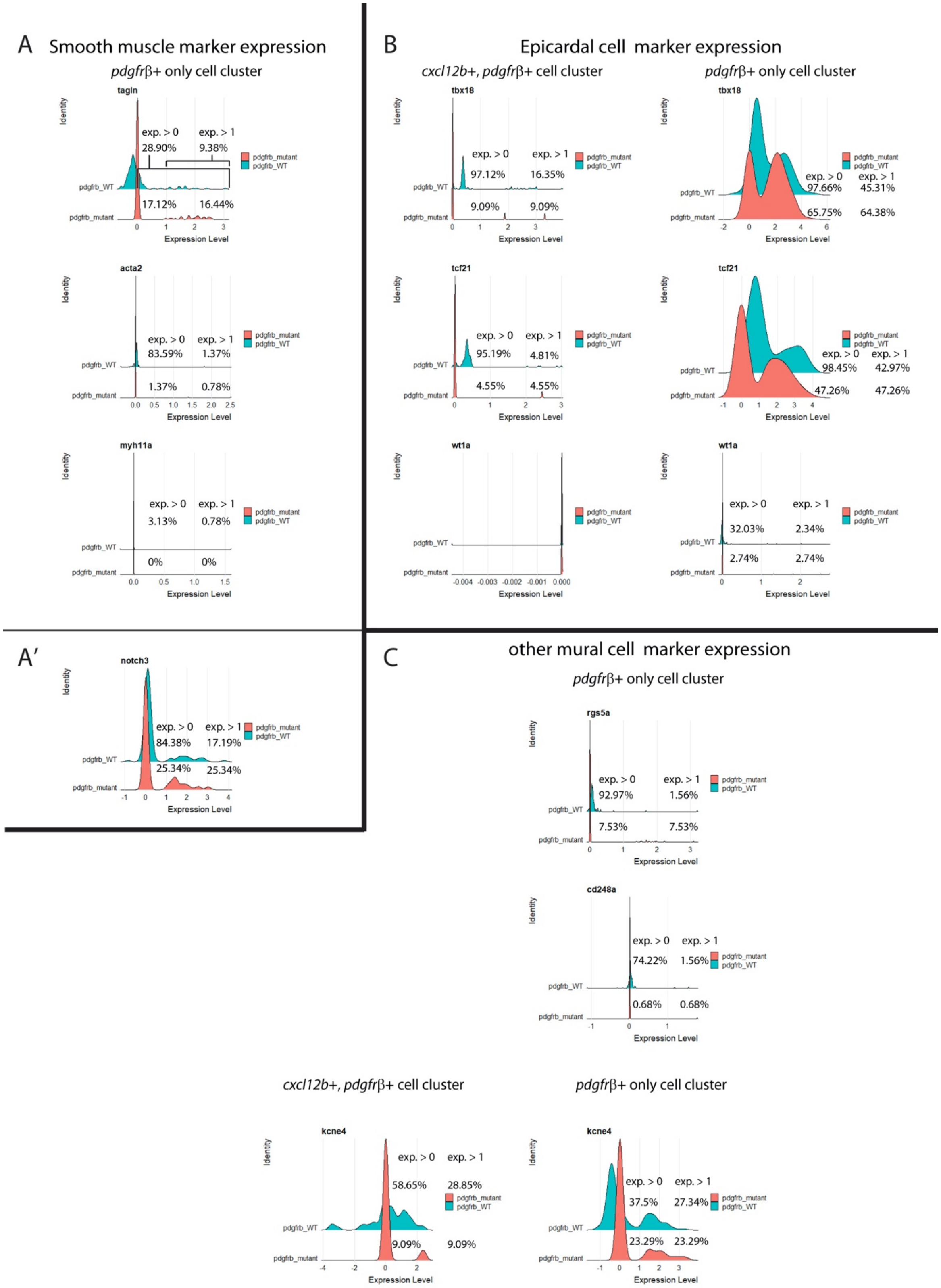
A Smooth muscle marker expression
pdgfrβ+ only cell cluster
tagln
exp. > 0 28.90%
exp. > 1 9.38%
17.12%
16.44%
acta2
exp. > 0 83.59%
exp. > 1 1.37%
1.37%
0.78%
myh11a
exp. > 0 3.13%
exp. > 1 0.78%
0%
0%
B Epicardal cell marker expression
cxcl12b+, pdgfrβ+ cell cluster
pdgfrβ+ only cell cluster
tbx18
exp. > 0 97.12%
exp. > 1 16.35%
9.09%
9.09%
exp. > 0 97.66%
exp. > 1 45.31%
65.75%
64.38%
tcf21
exp. > 0 95.19%
exp. > 1 4.81%
4.55%
4.55%
exp. > 0 98.45%
exp. > 1 42.97%
47.26%
47.26%
wt1a
exp. > 0 32.03%
exp. > 1 2.34%
2.74%
2.74%
A'
notch3
exp. > 0 84.38%
exp. > 1 17.19%
25.34%
25.34%
C other mural cell marker expression
pdgfrβ+ only cell cluster
rgs5a
exp. > 0 92.97%
exp. > 1 1.56%
7.53%
7.53%
cd248a
exp. > 0 74.22%
exp. > 1 1.56%
0.68%
0.68%
cxcl12b+, pdgfrβ+ cell cluster
pdgfrβ+ only cell cluster
kcne4
exp. > 0 58.65%
exp. > 1 28.85%
9.09%
9.09%
exp. > 0 37.5%
exp. > 1 27.34%
23.29%
23.29%
Identity
Expression Level
pdgfrb_WT
pdgfrb_mutant

**Supplementary Figure 4. Dynamic *pdgfrβ* expression during heart regeneration.**

(A) *pdgfrβ* expression during heart regeneration. Images of *Tg(pdgfrβ: Citrine; fli1a:DsRed)* fish hearts at 7, 14, 30 dpa. Lower panels: *pdgfrβ:citrine* single channel images. White dashed line: injuried area. White dotted line: *pdgfrβ* expressing area. White arrowhead, *pdgfrβ* expression in mural cells; yellow arrowhead, epicardial/non-mural cell *pdgfrβ* expression. (A') Quantification of *pdgfrβ*+ area as the percentage of the imaged heart area. Error bars, standard deviation of mean. One-way ANOVA (*$p < 0.05$). 1d, 3d, 30d; n = 3, 7d; n = 5, 10d, 14d; n = 4.

Tg(*pdgfr*β:*Citrine*; *fli1a:dsRed*)

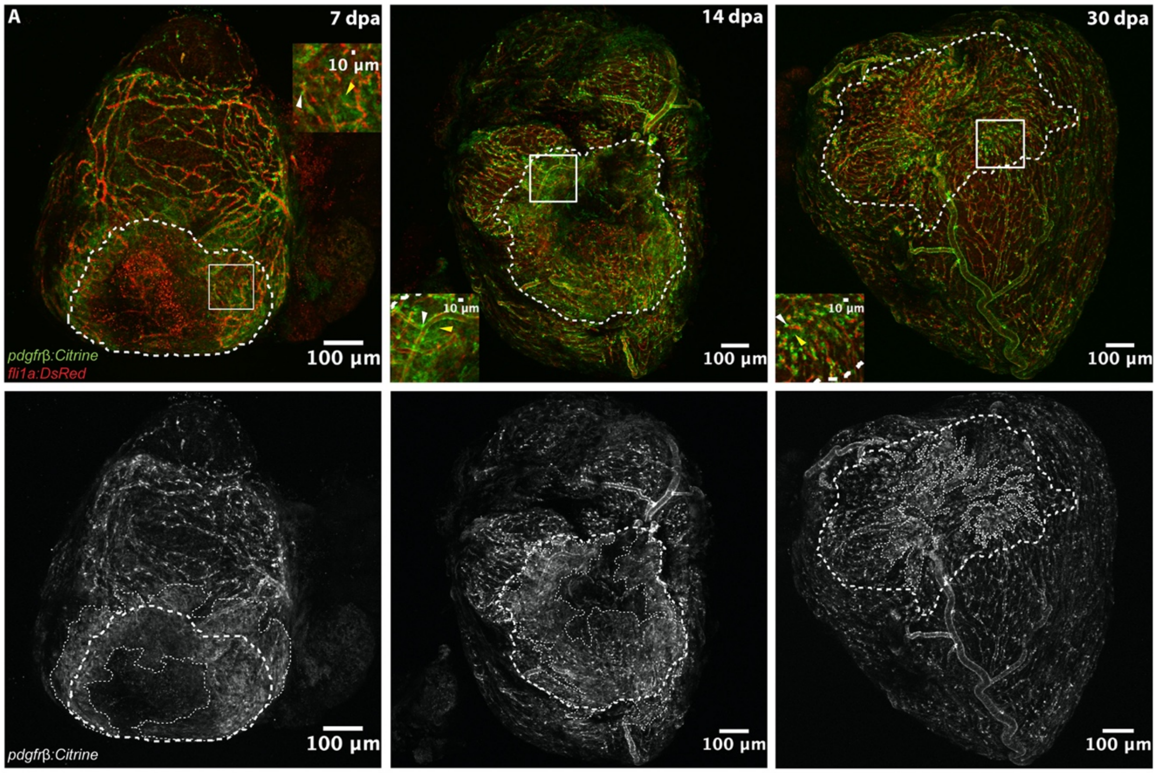

**Supplementary Figure 5.1. Identification and characterization of the epicardial cells for uninjured vs. injured hearts.** (A-A') *pdgfr$\beta$:EGFP* cells are FACS isolated from *Tg*(*pdgfr$\beta$:EGFP*; *cxcl12b:Citrine*) fish. (A) UMAP plot of all isolated EGFP+ cells (the uninjured and the 7 days post amputated samples integrated together), which form 15 clusters. (A') Feature plots for *pdgfr$\beta$* gene show the cells in the cluster 5, 6, 10, 12 express *pdgfr$\beta$*. (B) Cell type identification by marker gene expression. Violin plots are showing EGFP mRNA from the *pdgfr$\beta$:EGFP* reporter is mostly expressed in the cluster 0, 5, 6, 10, 11, 12, among which, cluster 5, 6, 10, 12 strongly express *pdgfr$\beta$*. Cluster 5, 6, 10 express epicardial marker genes (*tcf21*, *tbx18*, *wt1a*). Cluster 0, 11, 13 mostly expressed endothelial/endocardial cell markers (*fli1a*, *kdrl*). Cluster 1, 2, 3, 4 highly express cardiomyocyte marker *cmlc1*. Cluster 8 highly express red blood cell marker *hbba1*. Cluster 9, 14 express macrophage marker (*mpeg1.1*). Cluster 7 express T-cell marker (*lck*) and cluster 7, 9, 14 express lymphocyte marker (*lcp1*). (C) UMAP of all the isolated cells showing the cell clusters according to cell types. Cluster 5, 6, 10 (epicardial/EPDC/mural cells) and 12 (mural cells) cells (*pdgfr$\beta$* expressing clusters) are isolated and re-clustered into 5 subclusters.

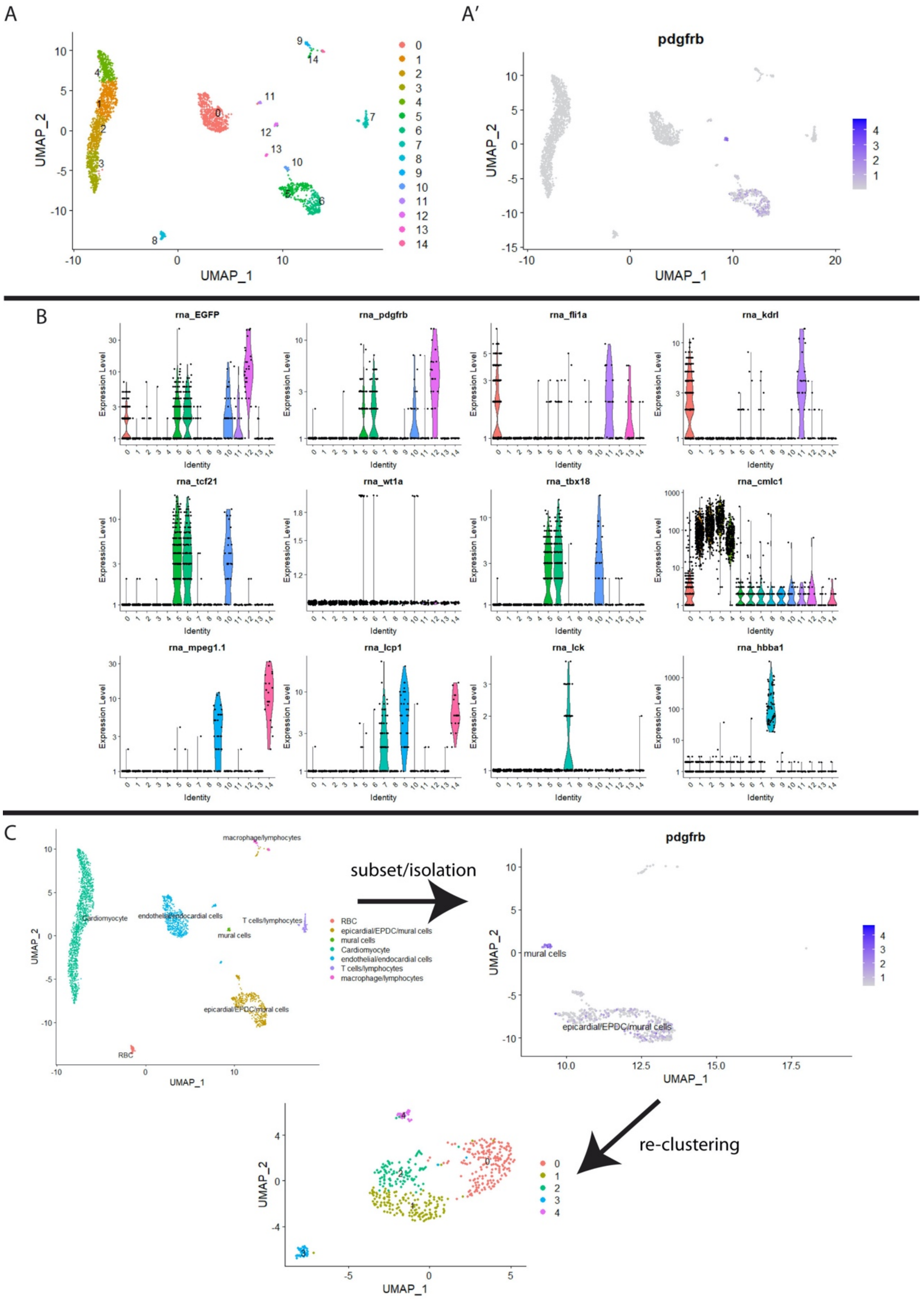
A
A′
pdgfrb
UMAP_1
UMAP_2
B
rna_EGFP
rna_pdgfrb
rna_fli1a
rna_kdrl
rna_tcf21
rna_wt1a
rna_tbx18
rna_cmlc1
rna_mpeg1.1
rna_lcp1
rna_lck
rna_hbba1
Expression Level
Identity
C
Cardiomyocyte
endothelial/endocardial cells
macrophage/lymphocytes
T cells/lymphocytes
mural cells
epicardial/EPDC/mural cells
RBC
subset/isolation
pdgfrb
mural cells
epicardial/EPDC/mural cells
re-clustering

**Supplementary Figure 5.2. Expression comparisons of the epicardial, epithelial to mesenchymal transition and pro-regenerative extracellular matrix genes between uninjured vs. injured (7 dpa) hearts.** (A) UMAP plot of re-clustered epicardial/EPDC cells and mural cells from *Tg*(*pdgfrβ:EGFP*; *cxcl12b:Citrine*) fish, 5-7 hearts in each of uninjured and injured (7dpa) conditions. (B-D) Violin plots of genes between *pdgfrβ*+cell in ctrl (uninjured) and *pdgfrβ*+cell in 7dpa (injured) hearts. (B) Violin plots for epicardial markers (*tbx18*, *wt1a*, *wt1b*). (C) Comparative feature plots for epithelial to mesenchymal transition markers (*snai1a*, *snai1b*, *snai2*, *twist1b*). (D) Violin plots for extracellular matrix genes (*fn1a*, *fn1b*, *postnb*).

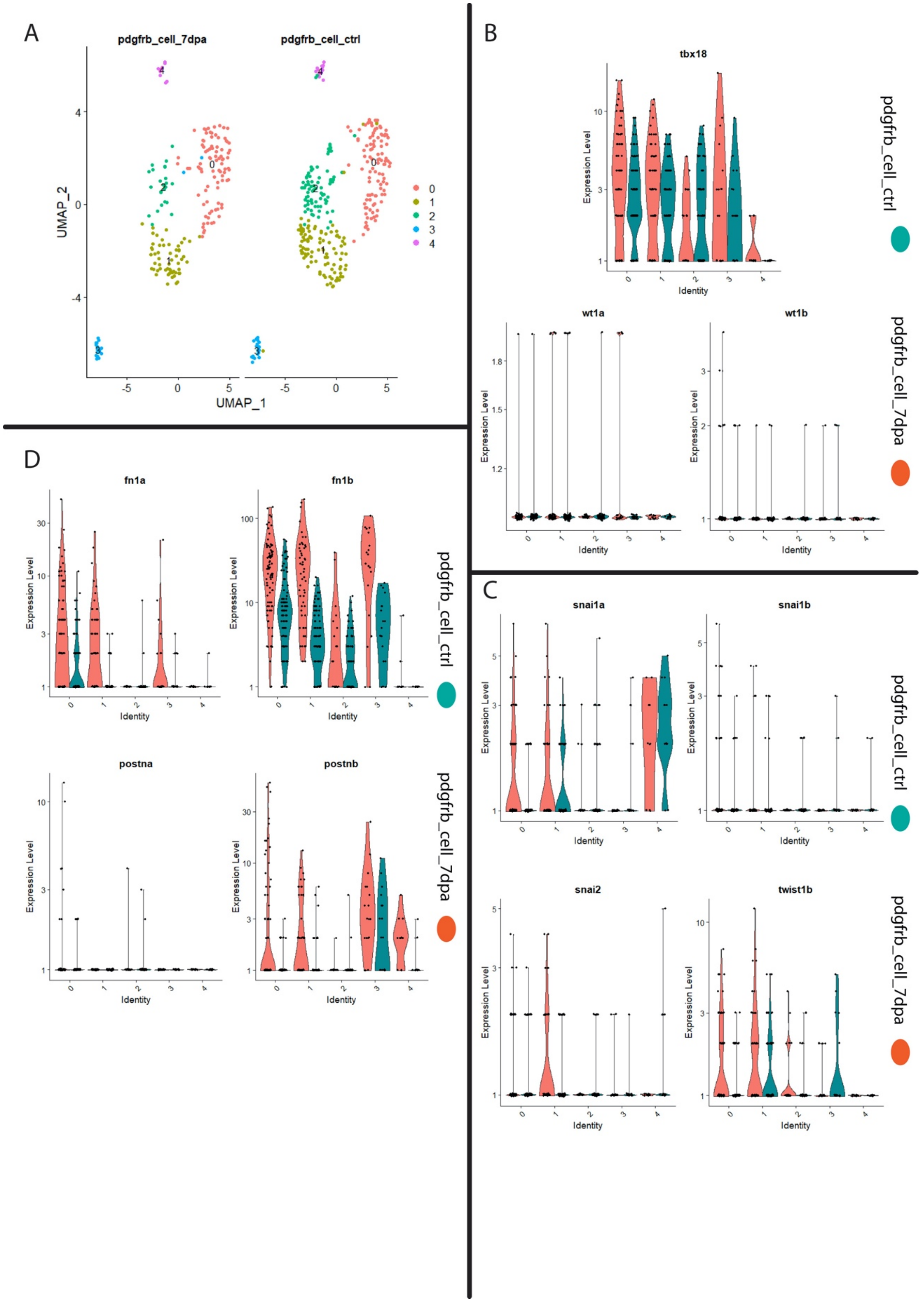

A
pdgfrb_cell_7dpa
pdgfrb_cell_ctrl
UMAP_2
UMAP_1
0
1
2
3
4
B
tbx18
wt1a
wt1b
Expression Level
Identity
pdgfrb_cell_ctrl
pdgfrb_cell_7dpa
C
snai1a
snai1b
snai2
twist1b
D
fn1a
fn1b
postna
postnb

**Supplementary Figure 6. Characterization of *mdka* in 7 dpa *pdgfrβ*+ cells and *mdka* mutants**

(A) *pdgfrβ:EGFP* cells and *fli1a:DsRed* cells were FACS sorted from 5, *Tg*(*pdgfrβ:EGFP; fli1a:DsRed*) fish and qRT-PCR was performed for *mdka* expression. The Y-axis shows expression fold changes of *mdka*, normalized to the housekeeping gene *rpl13a's* expression in uninjured vs. injured (7 days post-amputation of the ventricles' apical regions). (B) Representative images of in situ hybridization with the probe against *mdka* in injured (10 days post-amputation) wildtype fish vs. $mdka^{-/-}$ fish (n = 3). (C) Whole ventricles (n = 2) were collected from wild type uninjured hearts, injured (7 days post-amputation of apical regions of ventricles) hearts and $mdka^{-/-}$ injured (7 dpa) hearts. qRT-PCR was performed for *mdka*, and *pdgfrβ* expressions. The Y-axis shows expression fold changes normalized to the housekeeping gene *rpl13a's* expressions.

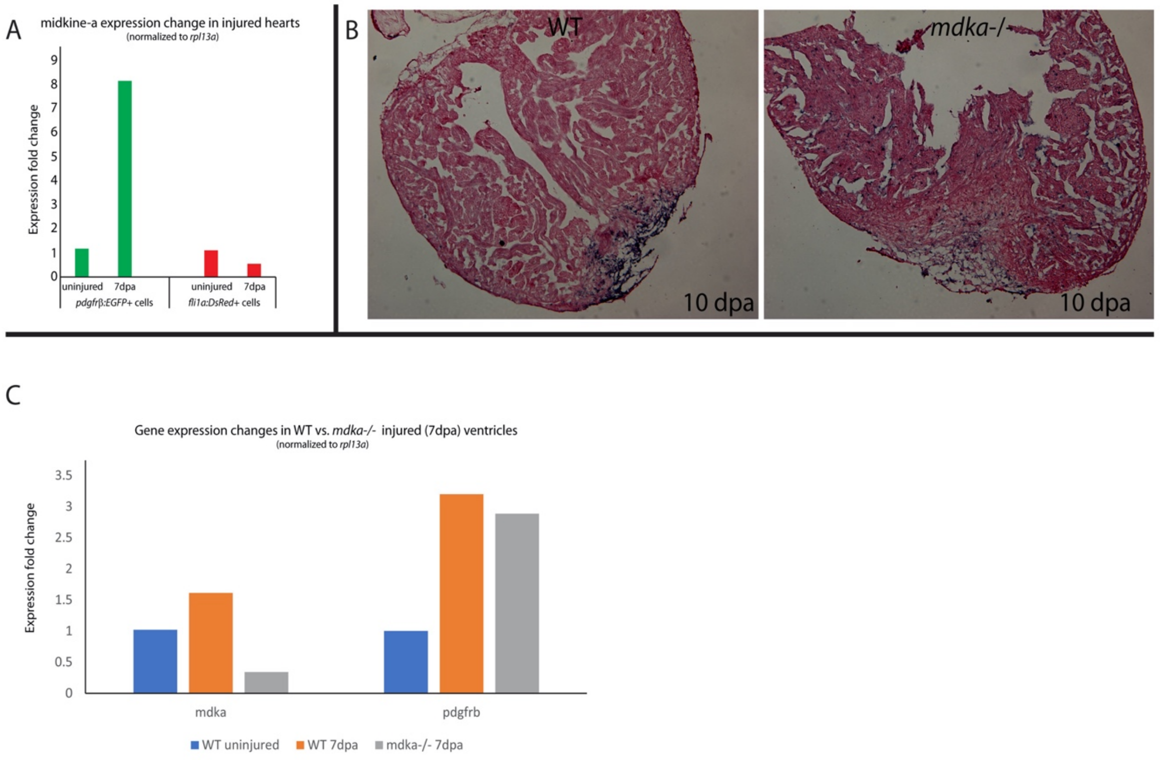
A
midkine-a expression change in injured hearts
(normalized to rpl13a)
Expression fold change
0
1
2
3
4
5
6
7
8
9
uninjured
7dpa
pdgfrβ:EGFP+ cells
uninjured
7dpa
fli1a:DsRed+ cells
B
WT
mdka-/-
10 dpa
10 dpa
C
Gene expression changes in WT vs. mdka-/- injured (7dpa) ventricles
(normalized to rpl13a)
Expression fold change
0
0.5
1
1.5
2
2.5
3
3.5
mdka
pdgfrb
WT uninjured
WT 7dpa
mdka-/- 7dpa

**Supplementary Movie**

Supplementary Movie 1

Movie from apex view showing behavior of two distinct *pdgfrfiEGFP* expression patterns, epicardial (diffuse) and mural (punctate), during regeneration at 5 to 6dpa. Time in minutes indicated on the bottom right.

Supplementary Movie 2

Movie from apex view showing *pdgfrfiEGFP (green) and fli1a:DsRed (red)* expression during regeneration 6 to 8dpa. Time in minutes indicated on the bottom right.

Supplementary Movie 3

Movie from lateral view showing *pdgfrfi:EGFP (red) and fli1a:DsRed (green)* expression during regeneration 10 to 12dpa. Time in minutes indicated on the bottom right.
